# Extensive sequencing of seven human genomes to characterize benchmark reference materials

**DOI:** 10.1101/026468

**Authors:** Justin M. Zook, David Catoe, Jennifer McDaniel, Lindsay Vang, Noah Spies, Arend Sidow, Ziming Weng, Yuling Liu, Chris Mason, Noah Alexander, Elizabeth Henaff, Feng Chen, Erich Jaeger, Ali Moshrefi, Khoa Pham, William Stedman, Tiffany Liang, Michael Saghbini, Zeljko Dzakula, Alex Hastie, Han Cao, Gintaras Deikus, Eric Schadt, Robert Sebra, Ali Bashir, Rebecca M. Truty, Christopher C. Chang, Natali Gulbahce, Keyan Zhao, Srinka Ghosh, Fiona Hyland, Yutao Fu, Mark Chaisson, Chunlin Xiao, Jonathan Trow, Stephen T. Sherry, Alexander W. Zaranek, Madeleine Ball, Jason Bobe, Preston Estep, George M. Church, Patrick Marks, Sofia Kyriazopoulou-Panagiotopoulou, Grace X.Y. Zheng, Michael Schnall-Levin, Heather S. Ordonez, Patrice A. Mudivarti, Kristina Giorda, Ying Sheng, Karoline Bjarnesdatter Rypdal, Marc Salit

**Affiliations:** National Institute of Standards and Technology, Gaithersburg, MD; Stanford University, Stanford, CA; Department of Physiology and Biophysics, the Feil Family Brain and Mind Research Institute, and HRH Prince Alwaleed Bin Talal Bin Abdulaziz Alsaud Institute for Computational Biomedicine, Weill Medical College, Cornell University, New York, NY 10065, USA.; Illumina Mission Bay, San Francisco, CA; BioNano Genomics, San Diego, CA; Department of Genetics and Genomic Sciences, Icahn School of Medicine at Mount Sinai, New York, NY; Complete Genomics Inc., Mountain View, CA, USA.; Thermo Fisher Scientific, South San Francisco, CA 94080; Genome Sciences, University of Washington, Seattle, WA; National Center for Biotechnology Information, National Library of Medicine, National Institutes of Health, 45 Center Drive, Bethesda, MD 20892, USA; PersonalGenomes.org, Boston, MA; Harvard Medical School, Boston, MA; 10X Genomics, Pleasanton, CA, United States, 94566; Department of Medical Genetics, Oslo University Hospital, Kirkeveien 166, Bygg 25, 0450, Oslo, Norway

## Abstract

The Genome in a Bottle Consortium, hosted by the National Institute of Standards and Technology (NIST) is creating reference materials and data for human genome sequencing, as well as methods for genome comparison and benchmarking. Here, we describe a large, diverse set of sequencing data for seven human genomes; five are current or candidate NIST Reference Materials. The pilot genome, NA12878, has been released as NIST RM 8398. We also describe data from two Personal Genome Project trios, one of Ashkenazim Jewish ancestry and one of Chinese ancestry. The data come from 12 technologies: BioNano Genomics, Complete Genomics paired-end and LFR, Ion Proton exome, Oxford Nanopore, Pacific Biosciences, SOLiD, 10X Genomics GemCode™ WGS, and Illumina exome and WGS paired-end, mate-pair, and synthetic long reads. Cell lines, DNA, and data from these individuals are publicly available. Therefore, we expect these data to be useful for revealing novel information about the human genome and improving sequencing technologies, SNP, indel, and structural variant calling, and de novo assembly.

## Background & Summary

Developing Reference Materials is a unique measurement science task, where significant resources can be expended to deeply characterize a small number of samples. Reference Materials act to calibrate, benchmark, or validate a measurement process. These samples are often the source of the scales on which we report our results (e.g., the Molar concentration of cholesterol), and they can be a physical realization of the SI units. Our ability to compare results between laboratories in most applications depends on Reference Materials.

There is a tradition of innovation in measurement science to characterize these high-impact samples.^1^ New technologies are used, rigorous experimental designs are employed, and exotic methods applied.^2,3^ In a virtuous cycle, existing methods are optimized and new methods are developed using reference materials as the benchmarks. Regulated applications depend on reference materials for quantitative, objective oversight; this opens new applications for a measurement technology, with great quality-of-life social benefit. With sequencing technologies and bioinformatics changing rapidly, whole genome reference materials and diverse data types like those presented here are a valuable resource for developing, improving, and assessing performance of these methods.

The National Institute of Standards and Technology (NIST)-hosted Genome in a Bottle Consortium is developing reference materials from well-characterized genomic DNA from 5 individuals (Figure 1). These reference materials are the first of their kind, and will play key roles in the translation of genome sequencing to widespread adoption and as validation tools in clinical practice. We previously characterized high-confidence SNP, indel, and homozygous reference genotypes,^4^ as well as large deletions and insertions.^5^ We plan to use similar methods as well as new methods to characterize these genomes using the data described in this work.

**Figure 1:**
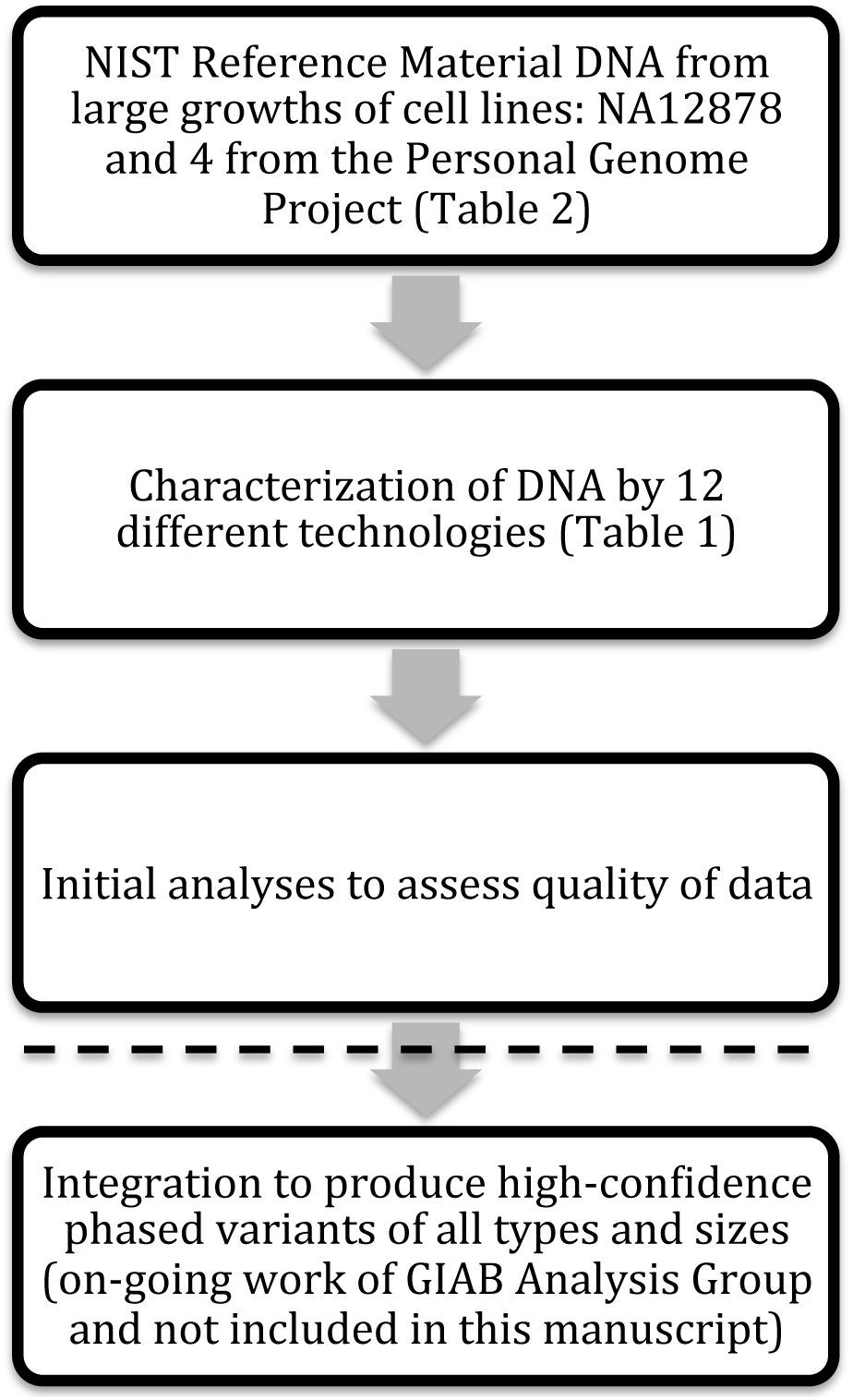
Overview of the study design, with data and analyses included in this manuscript above the dotted line and ongoing analyses of these data by the Genome in a Bottle Analysis Group below the dotted line.

The pilot genome (NIST RM 8398) is an oft-used genome: NA12878 from the CEPH Utah Reference Collection. In addition, genomes from two family trios (both Mother-Father-Son) have been selected from the Personal Genomes Project (PGP), one of Ashkenazi Jewish (AJ) ancestry and the other of Han Chinese ancestry. These genomes are available as cells or extracted DNA from the Coriell Institute for Medical Research and are or will be available as DNA as NIST Reference Materials. The NIST Reference Materials are extracted DNA from large, homogenized batches of cells prepared specially by Coriell to control for any batch effects. The samples from PGP are consented more broadly for many applications, including commercial redistribution. There are already three commercial products from the same cell lines from which the NIST Reference Material DNA is prepared: AcroMetrix^®^ Oncology Hotspot Control from Thermo Fisher Scientific, GIAB HDx^®^ Reference Standards from Horizon Diagnostics, and cell line DNA with synthetic DNA spike-ins from SeraCare Life Sciences.

The NIST Reference Material DNA has been characterized to an unprecedented degree. We have collected a large diverse set of data from 12 sequencing technologies and library preparation methods (Table 1). These data include high-depth paired-end short read whole genome sequencing (WGS) and whole exome sequencing (WES), long mate-pair WGS, pseudo long read (“read clouds”) WGS, long read WGS, genome mapping, and exome sequencing. For each dataset, we describe the library preparation and sequencing methods, the currently available data records, and technical validation. We expect these data to be complementary to each other, so that we can use them to characterize a broad spectrum of phased variants of all sizes in as much of the genome as possible. We invite anyone to join in this open, public effort to characterize these genomes; thus, here, we describe the measurement methods and data as a public resource.

**Table 1:** Summary of data available for GIAB samples

| Lab | Technology | Library Type | Date Produced | Mean Paired end distance | Mean read length | Mean coverage per individual for AJ trio | Mean coverage per individual for Chinese trio |
| --- | --- | --- | --- | --- | --- | --- | --- |
| NIST | Illumina HiSeq 2500 Rapid SBS | WGS paired end PCR-free | Jul to Sep 2014 | 565 bp | 2x250 (Chinese son), 2x148 (others) | 296.83x | 300x (son)<br>100x/parent |
| NIST/Stanford | Illumina HiSeq 2500 v3/v4 | WGS mate-pair | Dec 2014 to Mar 2015 | 6000 to 7000 bp | 100 | 13x to 14x | 14x to 17x |
| NIST/Illumina | Illumina HiSeq 2500 v4 | Synthetic Long Reads | Mar to Apr 2015 | N/A | 2x125 | 20x to 30x | 20x to 30x |
| Oslo University Hospital | Illumina HiSeq 2500 v4 | WES paired end | Oct 2015 | 202 bp | 2x125 | 192x to 225x | 181x (Son) |
| 10X Genomics | 10X GemCode WGS | WGS Linked-Reads | June to July 2015 | 200 to 220 bp | 2 x 98 | 25x (son)<br>24x (mother)<br>22x (father) | None |
| Complete Genomics | Complete Genomics | WGS paired end | Sep 2014 | 388 | 26 | 101x | 98x |
| Complete Genomics | Complete Genomics | WGS LFR | Dec 2014 | 278 | 26 | 100x | None |
| Thermo Fisher | Ion Proton | WES | Jun 2014 | N/A | 190 | 1020x | 1036x |
| NIST | SOLiD 5500xl Wildfire | WGS single end | Jan to Aug 2015 | N/A | 55 to 57 | 79x (Son) | 62x (Son) |
| BioNano Genomics | BioNano Genomics | WG optical mapping | Feb to Dec 2014 | N/A | 251kb (Chinese son), 195kb (AJ Son), 213kb (AJ Mother), 246kb (AJ Father) | 92x (Mother), 87x (Father), 112x (Son) | 57x (Son) |
| NIST/Mt. Sinai | PacBio P6-C4 (90%) & P5-C3 (10%) | WGS single end | Oct 2014 to Mar 2015 | N/A | 10-11kb N50 | 69x son; 30-32x/parent | None |
| Weill Cornell* | Oxford Nanopore | 2D reads | May 2015 | N/A | 5.8kb | 0.017x (son) | None |

\* Data described in <sup>14</sup>

## Methods

### Illumina paired end WGS

#### Library Preparation

For the AJ trio, Chinese son, and NA12878, libraries were prepared from 6 vials of the NIST Reference Material DNA for each individual. For the Chinese parents, a single library was prepared from genomic DNA from the Coriell Institute for Medical Research. For each Reference Material, 12 (or 14 for NA12878) libraries were prepared in parallel using the Illumina TruSeq (LT) DNA PCR-Free Sample Prep Kits (FC-121-3001). Two (or 3 for NA12878) libraries each were made from the first and last tubes in the lot, two libraries each were prepared from four samples pulled randomly from each quarter of the lot. This library design is intended for homogeneity analyses not presented here.

DNA concentrations were measured using a Qubit 2.0 fluorometer (Life Technologies). Genomic DNA (1.5 ug) was fragmented using a Covaris S2 focused ultrasonicator in micro TUBE AFA Fiber Pre-Slit Snap-Cap 6x16mm micro tubes and the Covaris MicroTUBE holder (covaris part numbers 520045 and 500114, respectively) under the following conditions for a target insert size of 550 base pairs. Duty cycle: 10%; Intensity: 2.0; Cycles Per Burst: 200; Duration: 45 seconds; Mode: Frequency Sweeping; Displayed Power: 9W; Temperature: 5.5° to 6°C. After Fragmentation, DNA was cleaned up using illumina Sample Purification Beads. End Repair was performed in 0.2 mL PCR tubes on an MJ research PTC-200 thermal cycler. The optional end repair control was not used. Size selection was done using a 96-well 0.8 mL plate (Fisher Scientific Part # AB-0859), a magnetic stand-96 (Ambion part # AM10027) and the Illumina sample purification beads according to the 550 bp insert protocol.

Adenylation of 3’ ends was done in 0.2 mL PCR tubes on an MJ Research PTC-200 thermal cycler. The optional A-Tailing control was not used. Ligation of indexed paired-end adapters was done in 0.2 mL PCR tubes using the DNA adapter tubes included in the Illumina TruSeq (LT) DNA PCR-Free Sample Prep Kit on an MJ Research PTC-200 thermal cycler. The optional ligation control was not used. The libraries were cleaned up in a 96-well 0.8 mL plate (Fisher Scientific Part # AB-0859) and a magnetic stand-96 (Ambion part # AM10027) using the Illumina sample purification beads. The final libraries were run on an Agilent 2100 Bioanalyzer HS-DNA chip to verify fragment size distribution. Final library concentration was measured via qPCR using the KAPA library quantification kit for Illumina sequencing platforms (KAPA part # KK4835). Libraries were then pooled based to the qPCR quantification data. The pool was intentionally made uneven so as to acquire greater sequence depth from the libraries made from the first and last tubes in each lot. The pools were adjusted between sequencing runs based on index balance.

For the Chinese son, DNA libraries were prepared in the same manner as they were for the Ashkenazim trio. The initial pool was made based on quantification measurements made using an Agilent 2100 Bioanalyzer, qPCR was not performed. This initial pool was sequenced on an Illumina MiSeq. The index balance obtained from the MiSeq run was used to adjust the pool for Sequencing on an Illumina HiSeq. The pool was intentionally made uneven so as to acquire greater sequence depth from the libraries made from the first and last tubes in each lot. The pools were adjusted between sequencing runs based on index balance.

#### Sequencing

For NA12878, the AJ Trio, and the Chinese parents, the pooled TruSeq libraries were run on an Illumina HiSeq 2500 in Rapid mode (v1) with 2x148 paired end reads. Pooled Libraries were initially loaded at a concentration of 10 pM. loading concentration was adjusted accordingly on subsequent runs to balance the libraries as well as possible.

For the Chinese son, the libraries were sequenced on an Illumina HiSeq 2500 in rapid mode (v2) with 2x250 paired end reads. Pooled Libraries were initially loaded at a concentration based on the information from the MiSeq run. Loading concentration was adjusted accordingly on subsequent runs to optimize cluster density.

The runs were designed to get approximately 300x total coverage of each of NA12878, the AJ Trio, and the Chinese son, and 100x coverage of each of the Chinese parents.

### Illumina mate-pair WGS

#### Library Preparation

Mate Pair libraries were generated using Nextera Mate Pair Sample Preparation Kit (Illumina, Cat# FC-132-1001). Briefly, 4 μg of high molecular weight genomic DNA from the NIST Reference Materials (or from Coriell for the Chinese parents) was fragmented to about 7 kb in a 400 mL tagmentation reaction containing 12 μL of Tagment Enzyme at 55°C for 30 minutes. The tagmented DNA fragments were purified with Zymo Genomic DNA Clean & Concentrator™ Kit (Zymo Research, Cat# D4010). The gap in the tagmented DNA was filled with a Strand Displacement Polymerase in a 200 μL strand displacement reaction at 20°C for 30 minutes. DNA was then purified with AMPure XP Beads (0.5x vol, Beckman Coulter, Cat# A63880) and size-selected by 0.6% agarose gel electrophoresis in 0.5x TBE buffer. The 6-9 kb fragments were excised from gel and DNA was recovered using a ZymocleanTM Large Fragment DNA Recovery Kit (Zymo Research, Cat# D4045). Up to 600 μg of DNA was then circulated overnight at 30°C with Circularization Ligase in a 300 μL reaction.

After overnight circularization, the uncirculated linear DNA was removed by Exonuclease digestion. Both DNA Ligase and Exonuclease were inactivated by heat treatment and the addition of Stop Ligation Buffer. Circularized DNA was then sheared to smaller sized fragments (300-1000 bp) using Covaris S2 with T6 (6x32 mm) glass tube (Covaris, Part# 520031 and 520042) under these conditions: Intensity of 8, Duty Cycle of 20%, Cycles Per Burst of 200, Time of 40 sec, Temperature of 6-8°C.

The sheared DNA fragments that contain the biotinylated junction adapter are mate pair fragments. These fragments were isolated by binding to Dynabeads M-280 Streptavidin Magnetic Beads (Invitrogen, Part# 112-05D) in Bead Bind Buffer. The unbiotinylated molecules in solution are unwanted genomic fragments that are removed through a series of washes. All downstream reactions were carried out on bead and beads were washed between successive reactions. The sheared DNA was first end-repaired to generate blunt ends followed by an A-Tailing reaction to add a single “A” nucleotide to the 3’ ends of the blunt fragments. Then the Illumina T-tailed indexing adapters were ligated to the A-tailed fragments.

The adapter-ligated fragments were PCR amplified [98°C/1 min, 11 cycles of (98°C/10 sec, 60°C/30 sec, 72°C/30s), 72°C/5 min, 4°C /hold] to generate the final library. The amplified library was purified using AMPure XP Beads (0.67x vol) and eluted in Resuspension Buffer. The size distribution of the library was determined by running a sample on an Agilent Technologies 2100 Bioanalyzer. Library concentration was measured by the Qubit dsDNA HS Assay Kit (Life Technologies, Cat# Q32851).

#### Sequencing

Pooled Mate-Pair libraries were sequenced on an Illumina HiSeq 2500 in Rapid mode (v1) with 2x101 bp paired-end reads. The loading concentration was 9.5 pM. This Initial run was for library QC purposes prior to running high throughput.

The Mate-Pair libraries were also sequenced on an Illumina HiSeq 2500 in high output mode (v4) with 2x125 bp paired-end reads. Libraries were sequenced on individual lanes (not pooled). The template loading concentration for each lane was adjusted based on the cluster density from the QC run. Two replicate flowcells were sequenced simultaneously, each with 6 lanes of mate-pair libraries.

### Illumina read clouds (synthetic long reads) WGS

#### Library Preparation

Synthetic long-read libraries were generated using the TruSeq^®^ Synthetic Long-Read DNA Library Prep Kit (Illumina, Cat# FC-126-1001). 500ng of DNA from the NIST Reference Materials (or from Coriell for the Chinese parents) was sheared, end-repaired, A-tailed, and adapters ligated before size-selecting 9-11 kb fragments according to the manufacturer’s protocol (Illumina Part # 15047264 Rev. B). Each resulting library was then diluted and aliquoted across a 384-well plate to limit the number of molecules to be amplified by PCR in each well. Amplified products were then tagmented and indexed by a second round of PCR (see referenced protocol for conditions) before pooling and concentrating all 384 wells for final product size selection and validation, again according to manufacturer’s instructions.

#### Sequencing

The synthetic long-read libraries for each genome were pooled and sequenced on an Illumina HiSeq 2500 in high output mode (v4) with 2x125 bp paired-end reads. Pooled libraries from each genome were loaded on individual lanes, with two lanes of each genome sequenced. The loading concentration for each lane was adjusted based on the cluster density of a previous (failed) run.

### Illumina paired end WES

#### Library Preparation

Captured exome DNA libraries for the Ashkenazim Jewish trio and the Chinese son were prepared from 4 vials of NIST Reference Material DNA. The library preparation was performed using Agilent SureSelect Automated Library Prep and Capture System Protocol and the Agilent SureSelect Target Enrichment System kit for 96 reactions (Agilent Technologies).

The DNA concentration for each sample was measured by Qubit 2.0 fluorometer (Life Technologies). The DNA samples were normalised with Low TE buffer to 4 ng/uL. 50 uL of each sample was fragmented in a 130 uL 96 MicroTUBE Plate with 96 microTUBE Foil Seal (part numbers 520078 and 520073 respectively) using the Covaris E-Series E220 focused ultrasonicator. The samples were sheared to an average of 250 bp, using the following instrument shearing settings: Average fragment size: 250 bp; Acousty Duty Factor: 10 %; PeakIncidence Power, W: 140; Cycles Per Burst: 200; Treatment Time: 160 seconds; Temperature: 6°C.

After fragmentation, the sheared DNA was purified using Agencourt AMPure XP purification beads, and the fragment size was controlled using the TapeStation 2200 with D1000 Reagents and ScreenTape. Following purification, the sample ends were modified for SureSelect target enrichment, through end-repair, 3’end adenylation, and adaptor ligation. The prepared DNA was purified after each each modification step. Sample purification, GA end-repair, A-tailing, and adaptor ligation were performed on the Bravo Agilent NGS Workstation robot in Nunc DeepWell plates. The adapter-ligated DNA fragments were captured and amplified in a 0.2 mL ABGENE 96-well PCR plate using the Applied Biosystems Veriti 96-well thermocycler. The thermocycler was programmed with the following settings: 98°C/3 min, 10 cycles of (98°C/80 sec, 65°C/30 sec, 72°C/1 min), 72°C/10 min, and 4°C/hold.

The libraries were cleaned up in a Nunc DeepWell plate using Agencourt AMPure XP beads on the Agilent NGS Workstation. To verify the DNA fragments has a size distribution between 200 and 400 bp, the libraries were measured on the TapeStation 2200 with D1000 Reagents and ScreenTape. The library concentration was measured using the Quant-iT dsDNA High-Sensitivity Assay Kit with the Victor3 1420 Multilabel counter. 750 ng of each prepped library sample was aliquoted for hybridization to the SureSelect Capture Library. Hybridization of the DNA libraries and the SureSelct Capture libraries was performed in a Nunc DeepWell plate on the Agilent NGS Workstation, and in a 0. 2 mL ABGENE 96-well PCR plate on the Applied Biosystems Veriti 96-well thermocycler. The thermocycler was programmed with a 95°C hybridization step for 5 min followed by a 65°C hold step. The hybridized libraries were captured with SureSelect Binding Buffer and purified using Dynabeads MyOne Streptavidin T1 bead suspension.

Addition of indexing tags to the SureSelect enriched captured libraries was performed through PCR based amplification, using the Agilent NGS Workstation and Applied Biosystems Veriti 96-well thermocycler. The thermocycler was programmed with the following settings: 98°C/2 min, 10 cycles of (98°C/30 sec, 57°C/30 sec, 72°C/1 min), 72°C/10 min, and 4°C/hold.

The amplified and indexed DNA libraries were purified with Agencourt AMPure XP beads. Quality control of the amplified captured libraries was done using the TapeStation 2200 High Sensitivity D1000 Reagents and ScreenTape to ensure a fragment size of 300 – 400 bp and a concentration of 10 nM. To estimate the cluster density and achive the final library concentration, qPCR was performed for each library using the KAPA library quantification kit for Illumina sequencing platforms. All four samples were prepared equally.

#### Sequencing

The libraries were pooled and diluted to a 2 nM pool (0,5 nM from each library), based on the qPCR measures. The library pool was sequenced on the Illumina HiSeq 2500 sequencing platform in high output run mode with 2x125bp paired-end reads.

### 10X Genomics GemCode™ Libraries for Illumina Sequencing

#### Genomic DNA Extraction

Genomic DNA was purified using a modified version of the MagAttract^®^ HMW DNA Kit (QIAGEN, Germantown, MD) from GM12878, GM24149, GM24143 and GM24385 cells (Coriell, Camden, New Jersey). Briefly, 1 x 10^6^ cells per extraction were pelleted and washed with PBS at RT. The Proteinase K and RNaseA digestion was incubated for 30 min at 25 ^o^C. Genomic DNA was purified using MagAttract^®^ Suspension G with Buffer MB, washed twice with Buffer MW1, and twice with Buffer PE. Finally the beads were rinsed twice with nuclease-free water for exactly 60 seconds. DNA was eluted with Buffer AE and quantified using the Qubit dsDNA HS Assay Kit (Thermo Fisher Scientific, Waltham, MA).

#### GemCode Whole Genome Library Preparation and Sequencing

Sample indexed and partition barcoded libraries were prepared using the GemCode kit (10X Genomics, Pleasanton, CA). 1.2 ng of DNA was used for GEM reactions where DNA fragments were massively partitioned into molecular reactors to extend the DNA and introduce specific 14-bp partition barcodes. GEM reactions were thermal cycled (95 ^o^C for 5 min; cycled 18X: 4 ^o^C for 30 sec, 45 ^o^C for 1 sec, 70 ^o^C for 20 sec, and 98 ^o^C for 30 sec; held at 4 ^o^C) and purified using the GemCode protocol. Purified DNA was sheared to 800-bp (M220, Covaris, Woburn, MA). Peak incident power: 75.0 W; duty factor: 5.0%; cycles per burst: 200; treatment time; 50 (s), temperature: 20.0 ^o^C; sample volume 50 ul. End repair, Adenylation tailing of 3’ ends, universal adapter ligation and sample indexing were performed according to the manufacture’s recommendations. Whole genome GemCode libraries were quantified by qPCR (KAPA Library Quantification Kit for Illumina^®^ platforms, Kapa Biosystems, Wilmington, MA). The NA12878 library was pooled with the NA24149 library and run on an Illumina HiSeq 2500 in Rapid mode (v1) with paired end 2x98-bp, 14-bp I5 and 8-bp I7 reads. For analysis the demultiplexed results from three flow cells were combined for a total of approximately 1.25 billion and approximately 810 million reads for NA12878 and NA24149, respectively. The NA24385 and NA24143 libraries were each run individually on a single high output mode (v4) lane for approximately 958 and approximately 900 million reads, respectively. Sequencing results were analyzed using the GemCode Long Ranger Software Suite

### Complete Genomics WGS

#### Library Preparation

Except for the Chinese parents, the NIST reference material was used as input. In the case of the Chinese trio, the full trio was sequenced from cells purchased from Coriell, and the son (GM24631) was also sequenced from the NIST reference material. Library prep followed the basic approach detailed previously,^6^ but with a two adapter library protocol (library version 2). Briefly, sequencing substrates were generated by means of genomic DNA fragmentation to a median fragment length of about 450 base pairs and recursive directional adapter insertion with an intermediate type IIS restriction enzyme digestion. The resulting circles were then replicated with *Phi29* polymerase (RCR)^7^ by synchronized synthesis to obtain hundreds of tandem copies of the sequencing substrate, referred to as DNA nanoballs (DNBs) which were adsorbed to silicon substrates with grid-patterned arrays to produce DNA nanoarrays.

#### Sequencing

High-accuracy cPAL sequencing chemistry (Version 2 sequencing) was used on automated sequencing machines to independently read up to 19 bases adjacent to each of the four anchor insertion sites, resulting in a total of 29-base mate-paired reads (58 bases per DNB). DNB intensity information is interpreted with the following steps: 1) background removal, 2) image registration, 3) intensity extraction. The intensity data from each field were then subjected to base calling, which involved four major steps: 1) crosstalk correction, 2) normalization, 3) base calling, and 4) raw base score computation.

### Complete Genomics LFR

#### Library Preparation

The LFR libraries were constructed as described in ^8^, except using the two adapter library protocol (library version 2) described above. Because this protocol requires cells as input, Coriell cells were used rather than the NIST reference material DNA. Briefly, controlled random enzymatic fragmenting is applied to 100–130 pg of high molecular mass (HMM) DNA that is physically separated into 384 distinct wells. The resulting fragments are then amplified and ligated to uniquely barcoded adapters. After combining the 384 wells and performing a restriction digestion, the second adapter is attached. The resulting substrate is converted to DNBs and adsorbed to silicon substrates with grid-patterned arrays to produce DNA nanoarrays.

#### Sequencing

Sequencing was performed as described above for regular Complete Genomics WGS, with the additional step of sequencing the well ID barcodes.

### Ion exome sequencing

#### Library Preparation

Exome libraries for 4 NIST Reference Materials, the AJ trio and Chinese son, were prepared using Ion AmpliSeq™ Exome RDY Kit, with a mean insert size of 215bp. Each sample was assigned a distinct barcode: IonXpress_020 for NA24385, IonXpress_022 for NA24149, IonXpress_024 for NA24143, and IonXpress_026 for NA24631. Each barcode library is diluted to 100pM. The libraries were emulsion-amplified individually and enriched using Ion OneTouch™ 2 System and Ion PI™ Template OT2 200 Kit v4. Outputs from 4 OneTouch runs for each sample were pooled together.

#### Sequencing

Each sample was sequenced on 4 Ion Proton™ instruments using Ion PI™ Sequencing 200 Kit v4. BaseCalling and alignment were performed on a Torrent Suite v4.2 server.

### SOLiD 5500xl Wildfire WGS

#### Fragment Library Preparation and Sequencing of AJ son and Chinese son

DNA sequencing on a Life Technologies 5500xl Wildfire was performed according to manufacturers protocols with noted modifications for each genome. A semi-automated library preparation process was first performed for the Chinese son Reference Material DNA. A modified manual library preparation was performed for the AJ son Reference Material DNA in an attempt to obtain smaller libraries for AJ son to maximize efficiency of colony formation on the 5500W. The two procedures used for each genome are detailed below.

#### Chinese son Semi-Automated Library Preparation

A semi-automated library preparation using the AB Library Builder System was used to prepare Chinese son libraries for sequencing on a 5500xl Wildfire. The workflow to produce 5500W DNA fragment libraries from Chinese son human genomic DNA (gDNA) was as follows (also in Supplementary Figure 1):

Shearing of gDNA was performed using the Covaris g-Tube (PN 520079) in conjunction with the Covaris S2 Focused Ultrasonicator. To obtain a uniform intermediate size distribution of approximately 10kb, 2.0 ug of gDNA was initially “pre-sheared” using a Covaris g-Tube in an Eppendorf 5424 centrifuge. The g-tubes were centrifuged twice at 4200 rpm for 60 seconds in each direction. Shearing was completed using the Covaris S2 per the User Guide “Fragment Library Preparation Using the AB Library Builder System: 5500 Series SOLiD Systems” (PN 4460965 Rev. A). Approximately 1.5 ug of “pre-sheared” gDNA was further sheared using the Covaris S2. Shearing was assessed on an Agilent 2100 Bioanalyzer High Sensitivity DNA Chip (PN 5067-4626) which showed a broad distribution of sheared material with peak at approximately 175 bp.

The Life Technologies AB Library Builder System was used to partially automate the library preparation process. End Repair, Size Selection, PolyA Tailing and Adaptor Ligation were performed on the AB Library Builder System to generate 5500 DNA fragment libraries. The Life Technologies Library Builder Fragment Core Kit for 5500 Genetic Analysis Systems (PN 4463763) and Beckman Coulter Agencourt AMPure XP Reagent (PN A263800) were used to prepare 5500 libraries on the AB Library Builder System. Adaptor amounts were calculated, per the Library Builder User Guide, based on input mass for a given sample. Library Preparation input mass ranged from 1.0-1.5 ug of sheared DNA depending on the given sample.

The AB Library Builder 5500 libraries then underwent manual nick translation and Wildfire library conversion to prepare libraries compatible for sequencing on a Life Technologies 5500xl Wildfire. Wildfire conversion was performed per the Quick Reference “5500 W Series Genetic Analysis Systems: Conversion of 5500 Library to 5500 W Library” (PN 4477188 Rev. B). Six cycles of amplification were performed in the conversion process.

Following an AmPure XP Reagent Cleanup the final 5500W DNA fragment libraries were run on the Sage Science BluePippin automated DNA size selection and collection system to further narrow the size distribution of the final libraries. A BluePippin DNA 2% Dye-Free Agarose gel cassette with V1 Marker (PN BDF2010) was used to capture DNA in a target range of 200-300 bps. All 5500W library for a given sample was loaded into the assigned well on cassette and run per the BluePippin 2% Agarose Gel Cassette Quick Guide. Upon completion of size selection 40-60 uL of size selected library was removed from the elution well and cleaned and concentrated using a 1.8X Agencourt AMPure XP (PN A263800) cleanup. Cleaned-size selected DNA was eluted in 32 uL of TE buffer. Size selection assessed using a Bioanalyzer High Sensitivity DNA Chip and showed the final Chinese son 5500W libraries with a size distribution of approximately 200-350 bps with peak at approximately 285 bps.

#### AJ son Modified Manual Library Preparation

A modified manual library preparation process for the AJ son was used to obtain appropriately sized libraries for sequencing on a 5500xl Wildfire. The workflow to produce 5500W DNA fragment libraries from AJ son human genomic DNA (gDNA) was as follows (also in Supplementary Figure 2):

**Figure 2:**
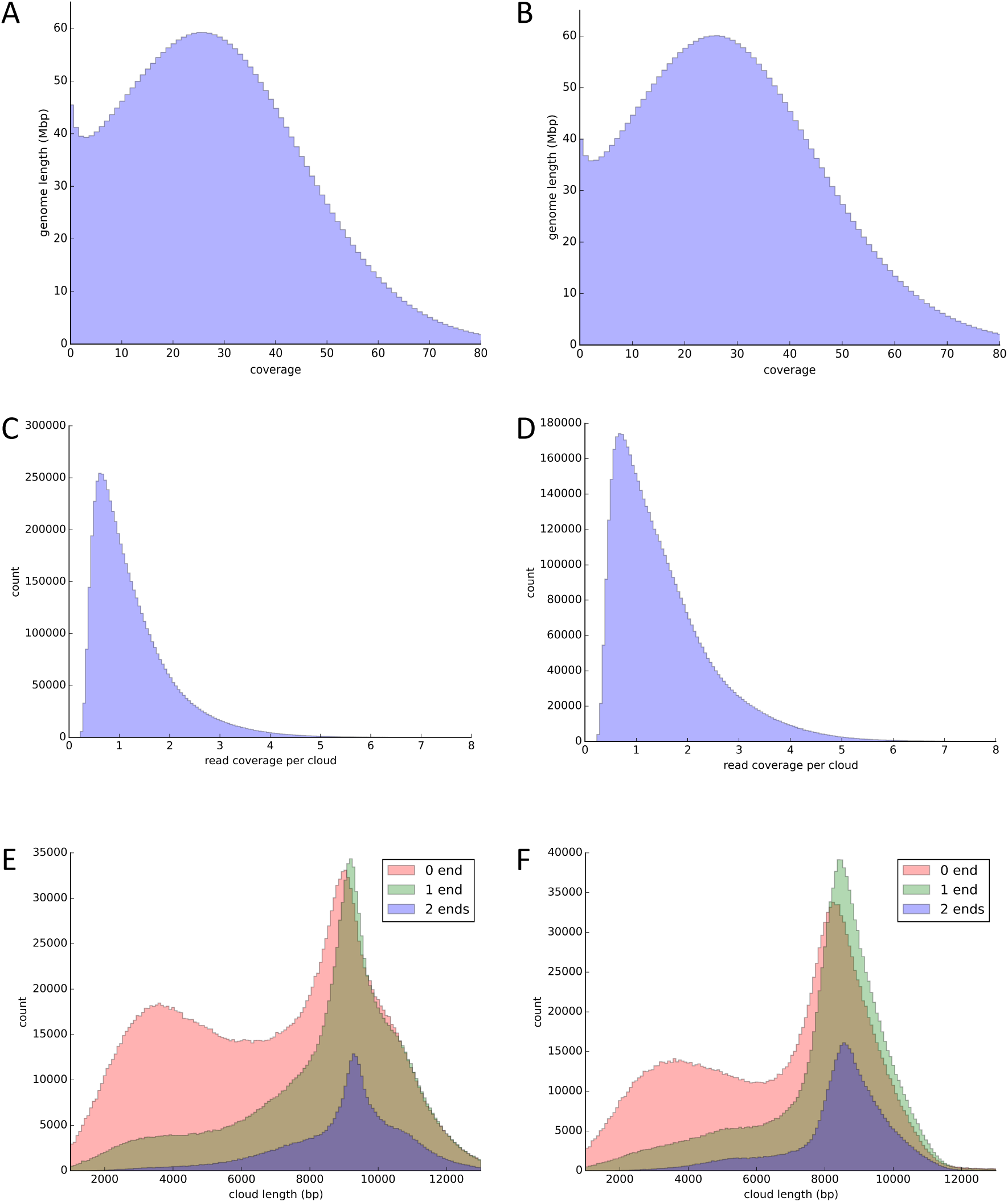
For HG002 (left - A, C, E) and HG003 (right - B, D, F), these are distributions of (A-B) coverage of the genome by short reads, (C-D) read coverage per cloud, and (E-F) cloud length for clouds with reads that contain markers at 0, 1, or 2 ends of the cloud.

Shearing of gDNA was performed using the Covaris g-Tube (PN 520079) in conjunction with the Covaris S2 Focused Ultrasonicator. To obtain a uniform intermediate size distribution of approximately 10kbp, 2.5 ug of gDNA was initially sheared using a Covaris g-Tube in an Eppendorf 5424 centrifuge. The g-tubes were centrifuged twice at 4200 rpm for 60 seconds in each direction. Shearing was completed using the Covaris S2 per the User Guide for “Fragment Library Preparation: 5500 Series SOLiD Systems” (PN 4460960 Rev. B). Approximately 2.0 ug of “pre-sheared” gDNA was sheared using the Covaris S2.

DNA fragment library preparation was performed using the total mass of sheared DNA (approximately 2.0 ug). Following the aforementioned 5500 Fragment Library Preparation guide, the 5500 SOLiD Fragment Library Core Kit (PN 4464412) was used to prepare 5500 libraries. The ends of the DNA fragments were repaired and DNA was cleaned and concentrated. Prior to size selection, the fragmented end-repaired DNA was assessed on an Agilent 2100 Bioanalyzer High Sensitivity DNA Chip (PN 5067-4626). Shearing resulted in a broad distribution with peak at approximately 175 bps.

To obtain a narrow fragment size distribution of DNA for AJ son 5500 library preparation, the DNA was run on the Sage Science BluePippin automated DNA size selection and collection system. A BluePippin DNA 3% Dye-Free Agarose gel cassette with Q2 Marker (PN BDF310) was used to capture DNA in a target range of 100-150 bps. Approximately 1.0-1.5 ug of end-repaired DNA was loaded into an appropriate well on the cassette and run per the BluePippin 3% Agarose Gel Cassette Quick Guide. Upon completion of size selection 40-60uL of size selected DNA was removed from the elution well and cleaned and concentrated using a 1.8X Agencourt AMPure XP (PN A263800) cleanup. Cleaned size-selected DNA was eluted in 32 uL of TE buffer. Size selection was again assessed using a Bioanalyzer High Sensitivity DNA Chip and showed a peak at approximately 128 bps.

Using reagents provided in the 5500 SOLiD Fragment Library Core Kit, dA Tailing, adaptor ligation and nick translation were performed per the 5500 Fragment Library Preparation Guide. Adaptor volumes were calculated using the mass calculated from the Bioanalyzer High Sensitivity Chip following the cleanup of the size-selected DNA. Two rounds of cleanup using Agencourt AMPure XP reagent were performed per the User Guide.

The completed 5500 libraries were then converted to 5500W libraries compatible for sequencing on a Life Technologies 5500xl Wildfire. Wildfire conversion was performed per the Quick Reference “5500 W Series Genetic Analysis Systems: Conversion of 5500 Library to 5500 W Library” (PN 4477188 Rev. B) utilizing reagents provided in the Life Technologies 5500W Conversion Primer Kit (PN 4478020) and Platinum PCR SuperMix (PN 11306-081). Six cycles of amplification were performed in the conversion process. Following completion of the conversion process, cleaned and concentrated libraries were assessed using a Bioanalyzer High Sensitivity DNA Chip and showed a peak at approximately 270 bps with a distribution from approximately 150-400 bps.

A second round of BluePippin size selection was performed to tighten the size distribution of the final 5500W library. The 5500W libraries were run on a DNA 2% Dye-Free Agarose gel cassette with V1 Marker to capture DNA in a target range of 200-300 bps. All DNA for a given sample was loaded into the assigned well on the cassette and run per the BluePippin 2% Agarose Gel Cassette Quick Guide. Upon completion of size selection 40-60uL of size selected DNA was removed from the elution well and cleaned and concentrated using a 1.8X Agencourt AMPure XP cleanup. Cleaned-size selected 5500W libraries were eluted in 32 uL of TE buffer. Size selection was assessed using a Bioanalyzer High Sensitivity DNA Chip and showed the final AJ son 5500W libraries with a peak at approximately 275 bps and a distribution from approximately 240-320 bps.

#### Sequencing of AJ son and Chinese son

A Life Technologies 5500xl Wildfire (5500W Genetic Analysis System) was used to sequence 5500W AJ son and Chinese son libraries using ICS software version 2.1. The User Guide “5500 W Series Genetic Analysis System (Americas)” (PN 4481746 Rev. B) was followed and used to prepare the samples and load a 5500W v2 FlowChip (PN 4475661). The Wildfire Template Amplification Protocol v6.1, located in the User Guide, was followed for template amplification. The 5500W FlowChip Prep Enzyme Kit (PN 4481058) and 5500W Template Amplification Reagents v2 (PN 4475663) were used to prepare FlowChips for “on-instrument” template amplification following template hybridization. 5500W library molar concentrations were calculated from the Bioanalyzer High Sensitivity chip following the final size selection of the 5500W libraries. These concentrations were used in calculation of FlowChip loading concentrations. Libraries were deposited into individual lanes at final concentrations of 100 to 250 pM. The library concentrations vary due to adjustments in subsequent instrument runs to increase colony density for a given library on the FlowChip.

5500W fragment libraries were sequencing with single-end 75 bp reads using the 5500W Forward SR 75 Reagent (PN 4475685). Two libraries were prepared and sequenced for each genome for a total of 24 lanes (4 FlowChips) per genome. This sequencing yielded approximately 72x coverage/ genome.

### Bionano Genomics genome maps

#### Library Preparation

Lymphoid-cell lines from the AJ Trio cell cultures obtained from Coriell Cell Repositories (GM24385, GM24143 and GM24149) were pelleted and washed with Life Technologies PBS (phosphate-buffered saline) at 1X concentration; the final cell pellet was re-suspended in cell suspension buffer using the Bio-Rad CHEF Mammalian Genomic DNA Plug Kit. Cells were then embedded in Bio-Rad CleanCut™ low melt Agarose and spread into a thin layer on a custom support in development. Cells were lysed using BioNano Genomics IrysPrep^®^ Lysis Buffer, protease treated with QIAGEN Puregene Proteinase K, followed by brief washing in Tris with 50mM EDTA and then washing in Tris with 1mM EDTA before RNase treatment with Qiagen Puregene RNase. DNA was then equilibrated in Tris with 50mM EDTA and incubated overnight at 4°C before extensive washing in Tris with 0.1mM EDTA followed by equilibration in New England BioLabs NEBuffer 3 at 1X concentration. Purified DNA in the thin layer agarose was labeled following the BioNano Genomics IrysPrep^®^ Reagent Kit protocol with adaptations for labeling in agarose. Briefly, 1.25 ug of DNA was digested with 0.7 units of New England BioLabs^®^ Nt.BspQI nicking endonuclease per μl of reaction volume in New England BioLabs NEBuffer 3 for 130 minutes at 37°C, then washed with Affymetrix TE Low EDTA Buffer, pH 8.0, followed by equilibration with New England BioLabs 1x ThermoPol^®^ Reaction Buffer. Nick-digested DNA was then incubated for 70 minutes at 50°C using BioNano Genomics IrysPrep^®^ Labeling mix and New England BioLabs Taq DNA Polymerase at a final concentration of 0.4U/μl. Nick-labeled DNA was then incubated for 40 minutes at 37°C using BioNano Genomics IrysPrep^®^ Repair mix and New England BioLabs^®^ Taq DNA Ligase at a final concentration of 1 U/μl. Labeled-repaired DNA was then recovered from the thin layer agarose by digesting with GELase™and counterstained with BioNano Genomics IrysPrep^®^ DNA Stain prior to data collection on the Irys system.

DNA was isolated from a lymphoid-cell cell culture of the Chinese son (GM24631) using the Bio-Rad CHEF Mammalian Genomic DNA Plug Kit protocol and lysed using BioNano Genomics IrysPrep^®^ Lysis Buffer and digested with QIAGEN Puregene Proteinase K. DNA was solubilized using GELase™ Agarose Gel-Digesting Preparation and drop-dialyzed before labeling using standard IrysPrep^®^ Reagent Kit protocols.

#### Mapping

Labeled and stained DNA samples were loaded into BioNano Genomics IrysChips^®^ and run on the BioNano Genomics Irys^®^ System imaging instrument. Data was collected for each sample until desired fold coverage of long molecules (>150 kb) was achieved. BioNano Genomics IrysView^®^ visualization and analysis software application was used to detect individual linearized DNA molecules using the Life Technologies YOYO^®^-1 Iodide in DMSO and determine the localization of labeled nick sites along each DNA molecule. BioNano Genomics IrysSolve™ analytical and assembly pipeline compiled the sets of single-molecule maps for each sample and were then used to build a full genome assembly.

### Pacific Biosciences

#### SMRTbell library preparation of AJ Trio gDNA

DNA library preparation and sequencing was performed according to the manufacturer’s instructions with noted modifications. Following the Pacific Biosciences Protocol, “20-kb Template Preparation Using Blue Pippin Size-Selection System”, library preparation was performed using the Pacific Biosciences SMRTbell Template Prep Kit 1.0 (PN # 100-259-100). In short, 10 μg of extracted, high-quality, genomic DNA from the NIST Reference Material DNA for the AJ trio, were used for library preparation. Genomic DNA extracts were verified with the Life Technologies Qubit 2.0 Fluorometer using the High Sensitivity dsDNA assay (PN# Q32851) to quantify the mass of doublestranded DNA present. After quantification, each sample was diluted to 150 μL, using kit provided EB, yielding a concentration of approximately 66 ng/μL. The 150 μL aliquots were individually pipetted into the top chambers of Covaris G-tube (PN# 520079) spin columns and sheared for 60 seconds at 4500 rpm using an Eppendorf 5424 benchtop centrifuge. Once complete, the spin columns were flipped after verifying that all DNA was now in the lower chamber. The columns were spun for another 60 seconds at 4500 rpm to further shear the DNA and place the aliquot back into the upper chamber. In some cases G-tubes were centrifuged 2-3 times, in both directions to ensure all volume had passed into the appropriate chamber. Shearing resulted in a approximately 20,000 bp DNA fragments verified using an Agilent Bioanlyzer DNA 12000 gel chip (PN# 5067-1508). The sheared DNA isolates were then purified using a 0.5X AMPure PB magnetic bead purification step (0.5X AMPure PB beads added, by volume, to each DNA sample, vortexed for 10 minutes at 2,000 rpm, followed by two washes with 70% alcohol and finally eluted in EB). This AMPure purification step assures removal of any small fragment and/or biological contaminant. The sheared DNA concentration was then measured using the Qubit High Sensitivity dsDNA assay. These values were used to calculate actual input mass for library preparation following shearing and purification. After purification, approximately 8 to 9 μg of each purified sheared sample went through the following library preparation process per this protocol (also in Supplementary Figure 3):

**Figure 3:**
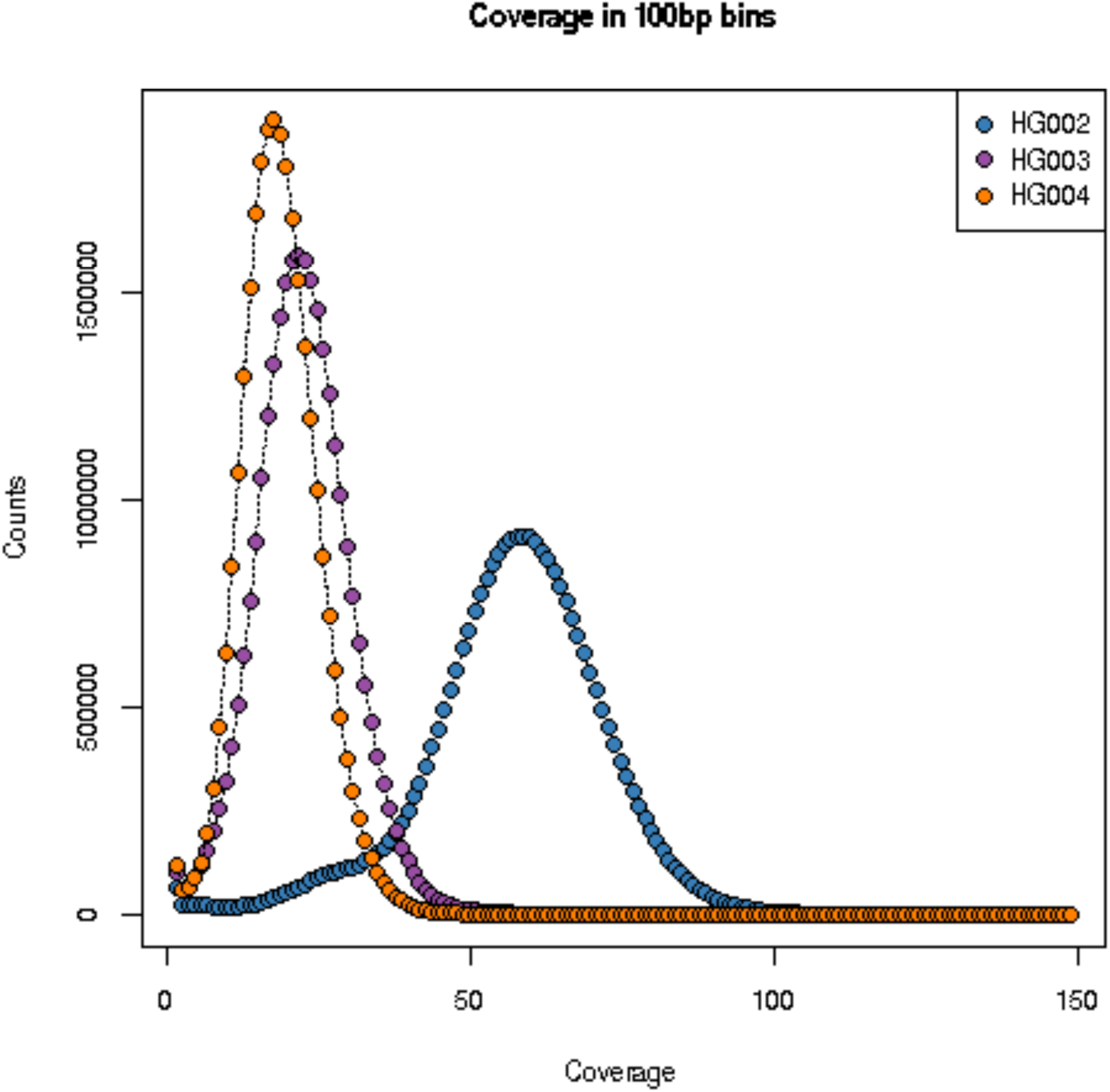
Histogram of coverage from PacBio for the AJ trio using a 100bp bin size

All library preparation reaction volumes were scaled to accommodate input mass for a given sample. Library size selection was performed using the Sage Science BluePippin 0.75% Agarose, Dye Free, PacBio approximately 20kb templates, S1 cassette (PN# PAC20KB). Size selections were run overnight to maximize recovered mass. Approximately 2-5 mg of prepared libraries were size selected using a 10 kb start and 50 kb end in “Range” mode. This selection is necessary to narrow the library distribution and maximize the SMRTbell sub-read length for the best *de novo* assembly possible. Without selection, smaller 2000 – 10,000 bp molecules dominate the zero-mode waveguide loading distribution, decreasing the sub-read length. Size-selection was confirmed using pre and post size selected DNA using an Agilent DNA 12000 chip. Final library mass was measured using the Qubit High Sensitivity dsDNA Assay. Approximately 15-20% of the initial gDNA input mass resulted after elution from the agarose cassette, which was enough yield to proceed to primer annealing and DNA sequencing on the PacBio RSII instrument. This entire library preparation and selection strategy was conducted 7, 2 and 2 times across AJ son, AJ father, and AJ mother respectively, to provide enough library for the duration of this project.

#### Sequencing AJ Trio on Pacific Biosciences RSII

Sequencing reflects the P6-C4 sequencing enzyme and chemistry, respectively. (Note that 10.3 % of the data was collected using the P5-C3 enzyme/chemistry prior to the release of the P6-C4 enzyme and chemistry.) Primer was annealed to the size-selected SMRTbell with the full-length libraries (80ºC for 2 minute 30 followed by decreasing the temperature by 0.1º/s to 25Cº). To prepare the polymerase-template complex, the SMRTbell template complex was then bound to the P6 enzyme using the Pacific Biosciences DNA Polymerase Binding Kit P6 v2 (PN# 100-372-700). A ratio of 10:1, polymerase to SMRTbell at 0.5 nM, was prepared and incubated for 4 hours at 30<u>°</u>C and then held at 4ºC until ready for magbead loading prior to sequencing. The Magnetic bead-loading step was conducted using the Pacific Biosciences MagBead Kit (PN# 100-133-600) at 4ºC for 60-minutes per manufacturer’s guidelines. The magbead-loaded, polymerase-bound, SMRTbell libraries were placed onto the RSII instrument at a sequencing concentration of 100 to 40 pM to optimize loading across various SMRTcells. Sequencing was performed using the C4 chemistry provided in the Pacific Biosciences DNA Sequence Bundle 4.0 (PN# 100-356-400). The RSII was then configured for at least 240-minute continuous sequencing runs.

### Oxford Nanopore

Methods to generate 2D reads from the Oxford Nanopore MinION from two different libraries are described in ^14^.

## Data Records

### Genomic samples

The genomes sequenced in this work (see Table 2) and their data are all publicly available both as EBV-immortalized B lymphoblastoid cell lines (from Coriell only) and as DNA (from Coriell and NIST). As described in the Methods, most data are from the NIST Reference Materials, unless the technology benefited from preparing longer DNA directly from cells. These human subjects are approved for “Public posting of personally identifying genetic information (PIGI)” by the Coriell and NIH/NIGMS IRBs, and this study was approved by NIST and by the Coriell/NIGMS IRB.

**Table 2:** Genomes currently being characterized by the Genome in a Bottle Consortium (NCBI BioProject PRJNA200694)

| Genome | Coriell cell line ID | NIST ID | NIST RM # | NCBI BioSample | PGP ID |
| --- | --- | --- | --- | --- | --- |
| CEPH Mother/Daughter | GM12878 | HG001 | RM8398 | SAMN03492678 | Not PGP |
| AJ Son | GM24385 | HG002 | RM8391* (son)/RM8392* (trio) | SAMN03283347 | huAA53E0 |
| AJ Father | GM24149 | HG003 | RM8392* (trio) | SAMN03283345 | hu6E4515 |
| AJ Mother | GM24143 | HG004 | RM8392* (trio) | SAMN03283346 | hu8E87A9 |
| Chinese Son | GM24631 | HG005 | RM8393* | SAMN03283350 | hu91BD69 |
| Chinese Father | GM24694 | N/A** | N/A** | SAMN03283348 | huCA017E |
| Chinese Mother | GM24695 | N/A** | N/A** | SAMN03283349 | hu38168C |

\*Not currently available. Planned release as NIST RMs in early 2016
\*\*NIST Reference Materials are not planned for the Chinese parents, but cells and DNA are available from Coriell.

### Illumina paired end WGS

#### NA12878

Approximately 300x 148bp x 148bp Illumina paired end WGS data from NA12878 is in the NCBI SRA SRX1049768 to SRX1049855 [Data Citation 1].

#### AJ Trio

148x148bp HiSeq sequencing and analyses of two 40x to 50x runs from each member of the Ashkenazim trio using the BWA-GATK pipeline on Basespace. Raw data is available in the SRA: SRX847862 to SRX848317 [Data Citation 2]. The BAM, VCF, and fastq files have been uploaded to:

ftp://ftp-trace.ncbi.nlm.nih.gov/giab/ftp/data/AshkenazimTrio/HG002_NA24385_son/NIST_HiSeq_HG002_Homogeneity-10953946/

ftp://ftp-trace.ncbi.nlm.nih.gov/giab/ftp/data/AshkenazimTrio/HG003_NA24149_father/NIST_HiSeq_HG003_Homogeneity-12389378/

ftp://ftp-trace.ncbi.nlm.nih.gov/giab/ftp/data/AshkenazimTrio/HG004_NA24143_mother/NIST_HiSeq_HG004_Homogeneity-14572558/

#### Chinese Trio

Data from the Chinese trio is in the NCBI SRA SRX1388368 to SRX1388459 [Data Citation 3]. Fastq files for 300x sequencing of Chinese son, as well as approximately 45x bam files generated from each flow cells are located here:

ftp://ftp-trace.ncbi.nlm.nih.gov/giab/ftp/data/ChineseTrio/HG005_NA24631_son/HG005_NA24631_son_HiSeq_300x

Fastq files for 100x sequencing of the Chinese parents, as well as approximately 100x bam files generated for each genome are located here:

ftp://ftp-trace.ncbi.nlm.nih.gov/giab/ftp/data/ChineseTrio/HG006_NA24694-huCA017E_father/NA24694_Father_HiSeq100x

ftp://ftp-trace.ncbi.nlm.nih.gov/giab/ftp/data/ChineseTrio/HG007_NA24695-hu38168_mother/NA24695_Mother_HiSeq100x

## Illumina mate-pair sequencing

Illumina mate-pair data are available at the NCBI SRA SRX1388732 to SRX1388743 [Data Citation 4], and as bam files in the NIST_Stanford_Illumina_6kb_matepair directory for each genome:

ftp://ftp-trace.ncbi.nlm.nih.gov/giab/ftp/data/AshkenazimTrio/HG002_NA24385_son/NIST_Stanford_Illumina_6kb_matepair/

ftp://ftp-trace.ncbi.nlm.nih.gov/giab/ftp/data/AshkenazimTrio/HG003_NA24149_father/NIST_Stanford_Illumina_6kb_matepair/

ftp://ftp-trace.ncbi.nlm.nih.gov/giab/ftp/data/AshkenazimTrio/HG004_NA24143_mother/NIST_Stanford_Illumina_6kb_matepair/

ftp://ftp-trace.ncbi.nlm.nih.gov/giab/ftp/data/ChineseTrio/HG005_NA24631_son/NIST_Stanford_Illumina_6kb_matepair/

ftp://ftp-trace.ncbi.nlm.nih.gov/giab/ftp/data/ChineseTrio/HG006_NA24694-huCA017E_father/NIST_Stanford_Illumina_6kb_matepair/

ftp://ftp-trace.ncbi.nlm.nih.gov/giab/ftp/data/ChineseTrio/HG007_NA24695-hu38168_mother/NIST_Stanford_Illumina_6kb_matepair/

### Illumina read clouds (synthetic long reads)

Illumina read cloud data are available as fastq’s (for the AJ trio and Chinese trio) and as bam files (currently only for the AJ son and father) in the NIST_Stanford_Moleculo directory for each genome:

ftp://ftp-trace.ncbi.nlm.nih.gov/giab/ftp/data/AshkenazimTrio/HG002_NA24385_son/NIST_Stanford_Moleculo/

ftp://ftp-trace.ncbi.nlm.nih.gov/giab/ftp/data/AshkenazimTrio/HG003_NA24149_father/NIST_Stanford_Moleculo/

ftp://ftp-trace.ncbi.nlm.nih.gov/giab/ftp/data/AshkenazimTrio/HG004_NA24143_mother/NIST_Stanford_Moleculo/

ftp://ftp-trace.ncbi.nlm.nih.gov/giab/ftp/data/ChineseTrio/HG005_NA24631_son/NIST_Stanford_Moleculo/

ftp://ftp-trace.ncbi.nlm.nih.gov/giab/ftp/data/ChineseTrio/HG006_NA24694-huCA017E_father/NIST_Stanford_Moleculo/

ftp://ftp-trace.ncbi.nlm.nih.gov/giab/ftp/data/ChineseTrio/HG007_NA24695-hu38168_mother/NIST_Stanford_Moleculo/

### Illumina paired end WES

Illumina paired-end WES data for AJ Trio and the Chinese son are available. The sequencing data were aligned by bwa mem (Li 2013) against b37 human decoy reference genome. The alignments were sorted and PCR duplicates were marked by Picard (http://picard.S0Urcef0rge.net). For AJ trios, a joint variant calling was performed by GATK (McKenna et al., 2010) HaplotypeCaller on all three samples. For the Chinese son, both single sample variant calling (a VCF file) and the first step in cohort analysis (a gVCF file) were performed by GATK HaplotypeCaller. All variants in VCF files were quality filtered by standard GATK SNP variant quality score recalibration and indel hard filtration according to GATK Best Practices recommendations (DePristo et al., 2011; Van der Auwera et al.,2013). Raw data (fastq files) are available in the SRA: SRP047086 [Data Citation 5]. The BAM and VCF files have been uploaded to:

BAM files and their index files:

ftp://ftp-trace.ncbi.nlm.nih.gov/giab/ftp/data/AshkenazimTrio/HG002_NA24385_son/OsloUniversityHospital_Exome

ftp://ftp-trace.ncbi.nlm.nih.gov/giab/ftp/data/AshkenazimTrio/HG003_NA24149_father/OsloUniversityHospital_Exome

ftp://ftp-trace.ncbi.nlm.nih.gov/giab/ftp/data/AshkenazimTrio/HG004_NA24143mother/OsloUniversityHospital_Exome

ftp://ftp-trace.ncbi.nlm.nih.gov/giab/ftp/data/ChinesTrio/HG005_NA24631_son/OsloUniversityHospital_Exome

Joint Variant Calling file for AJ trio:

ftp://ftp-trace.ncbi.nlm.nih.gov/giab/ftp/data/AshkenazimTrio/analvsis/OsloUniversityHospital_Exome_GATK_jointVC_11242015

Single sample variant calling file for the Chinese son:

ftp://ftp-trace.ncbi.nlm.nih.gov/giab/ftp/data/ChineseTrio/analysis/OsloUniversityHospital_Exome_GATK_jointVC_11242015

gVCF for the Chinese son:

ftp://ftp-trace.ncbi.nlm.nih.gov/giab/ftp/data/ChineseTrio/analysis/OsloUniversityHospital_Exome_GATK_jointVC_11242015

### 10X Genomics GemCode™ Libraries for Illumina Sequencing

10X Genomics data was generated with cell lines acquired from Coriell. Aligned reads with barcode and phasing information are provided in BAM format for each sample. VCF files with small variants are also provided for each sample, and SV calls are provided for NA12878 and the AJ son. See http://software.10xgenomics.com/ for detailed information on file formats. 10X Genomics data are available at http://software.10xgenomics.com/giab2015

The same data are available at the NCBI SRA SRX1392293 to SRX1392296 [Data Citation 6] and on the GIAB FTP:

ftp://ftp-trace.ncbi.nlm.nih.gov/giab/ftp/data/AshkenazimTrio/HG002_NA24385_son/10XGenomics

ftp://ftp-trace.ncbi.nlm.nih.gov/giab/ftp/data/AshkenazimTrio/HG003_NA24149_father/10XGenomics

ftp://ftp-trace.ncbi.nlm.nih.gov/giab/ftp/data/AshkenazimTrio/HG004_NA24143_mother/10XGenomics

### Regular Complete Genomics WGS

The Complete Genomics files are available on the NCBI SRA [Data Citation 7] and [Data Citation 8], and on the GIAB FTP site as summarized in Table 3. For GM24631, the son of the Chinese trio, data is available for both the NIST reference material and cells sourced from Coriell. Data is available for the Chinese parents (GM24695 and GM24694) from cells sourced from Coriell. All other data is from NIST reference materials.

**Table 3:** Complete Genomics data available at ftp://ftp-trace.ncbi.nlm.nih.gov/giab/ftp/

| Genome | File types | Location |
| --- | --- | --- |
| HG-001 | VCF | /data/NA12878/analysis/variant_calls/COMPLETE/Library2/ |
| HG-001 | BAM | /data/NA12878/CompleteGenomics_normal_RMDNA/alignment/Library2/ |
| HG-001 | Full package | /technical/complete_data/HG001_NA12878_normalCG/GS000025639-ASM/EXP/GS02256-DNA_A01/ |
| Ashekenazi trio | Multi-sample VCF | /data/AshkenazimTrio/analysis/CompleteGenomics_RefMaterial_SmallVariants_CGAtools_08082014/AshkenaziTrio_RefMaterial_SmallVariants.vcf |
| GM24385 | BAM, VCF, reference scores | /data/AshkenazimTrio/HG002_NA24385_son/CompleteGenomics_normal_RMDNA/son_NA24385_GS000037263-ASM/ |
| GM24385 | Full package | /technical/complete_data/trios/native_format_data/GS000043892-DID/GS000037263-ASM/ |
| GM24143 | BAM, VCF, reference scores | /data/AshkenazimTrio/HG004_NA24143_mother/CompleteGenomics_normal_RMDNA/mom_NA24143_GS000037262-ASM/ |
| GM24143 | Full package | /technical/complete_data/trios/native_format_data/GS000043891-DID/GS000037262-ASM/ |
| GM24149 | BAM, VCF, reference scores | /data/AshkenazimTrio/HG003_NA24149_father/CompleteGenomics_normal_RMDNA/dad_NA24149_GS000037264-ASM/ |
| GM24149 | Full package | /technical/complete_data/trios/native_format_data/GS000043893-DID/GS000037264-ASM/ |
| Han trio (Coriell cells) | Multi-sample VCF | /data/ChineseTrio/analysis/CompleteGenomics_HanTrio_ExtractedFromCoriellCells_SmallVariants_CGAtools_08082014/HanTrio_ExtractedFromCoriellCells_SmallVariants.vcf |
| GM24631 | BAM, VCF, reference scores | /data/ChineseTrio/HG005_NA24631_son/CompleteGenomics_normal_RMDNA/son_NA24631_GS000037265-ASM/ |
| GM24631 | Full package | /technical/complete_data/trios/native_format_data/GS000043894-DID/GS000037265-ASM/ |
| GM24631 (cells) | BAM, VCF, reference scores | /data/ChineseTrio/HG005_NA24631_son/CompleteGenomics_normal_cellsDNA/son_NA24631_GS000037475-ASM/ |
| GM24631 (cells) | Full package | /technical/complete_data/trios/native_format_data/GS000044335-DID/GS000037475-ASM/ |
| GM24695 (cells) | BAM, VCF, reference scores | /data/ChineseTrio/HG007_NA24695-hu38168_mother/CompleteGenomics_normal_cellsDNA/mom_NA24695_GS000037477-ASM/ |
| GM24695 (cells) | Full package | /technical/complete_data/trios/native_format_data/GS000044337-DID/GS000037477-ASM/ |
| GM24694 (cells) | BAM, VCF, reference scores | /data/ChineseTrio/HG006_NA24694-huCA017E_father/CompleteGenomics_normal_cellsDNA/dad_NA24694_GS000037476-ASM/ |
| GM24694<br>(cells) | Full package | /technical/complete_data/trios/native_format_data/GS000044336-DID/GS000037476-ASM/ |

Directory structures and file formats for the “Full package”, as well as the other supplementary files discussed below, are described in http://www.completegenomics.com/documents/DataFileFormats_Standard_Pipeline_2.5.pdf. For both the Chinese and Askenazi trios, a multisample VCF including only small variants was generated from masterVar files using the CGA tools mkvcf program described in http://cgatools.sourceforge.net/docs/1.8.0/cgatools-user-guide.pdf. All other VCF files contain small variants, CNVs, SVs and MEIs. Note that for CNVs and SVs, more complete information is available in the ASM/CNV and ASM/SV directories of the full package. For CNVs, VCF files contain the information sourced from the cvnDetails files but do not provide information on any segmentation of the genome into ploidy or coverage levels. For SVs, VCF files contain information sourced from the allJunctionsBeta and highConfidenceJunctionsBeta, but information from the allSvEventsBeta and highConfidenceSvEventsBeta files is not included.

BAM files are provided in order to provide evidence of variants called. However, it is not appropriate to remap and recall variants based on these BAM files as proper re-mapping of reads should take into account the gapped read structure. The *_mapping_sorted_header.bam files include the initial mappings of all reads. They were generated with the map2sam program from CGA Tools with the --mate-sv-candidates and --add-unmapped-mate-info parameters. Inconsistent mappings are normally converted as single arm mappings with no mate information provided, but with the --mate-sv-candidates option map2sam will mate unique single arm mappings in SAM including those on different stands and chromosomes. The tag XS:i:1 is used to distinguish these ‘ ‘ artificially ‘ ‘ mated records. The MAPQ provided for these records is a single arm mapping weight. The --add-unmapped-mate-info parameter generates mate sequences and score tags for inconsistent mappings. In the subsequent local de novo assembly (LDN) stage of genome assembly, reads can be re-mapped, added or removed from the assembly within the region undergoing LDN. The reads and the mappings that support variant calls after LDN is complete are provided in the evidence files. EvidenceDnbs* bam files are generated with our evidence2sam tool from CGA Tools (http://cgatools.sourceforge.net/docs/1.8.0/cgatools-user-guide.pdf). A detailed description of the data file can be found in the, “Representation of the Complete Genomics Data in SAM Output Format” appendix of the CGA Tools User Guide (http://cgatools.sourceforge.net/docs/1.8.0/cgatools-user-guide.pdf). They contain the reads and mappings that support one of the called alleles by at least 2 dB over the other called allele. This means they will not contain reads and mappings that do not support either of the called alleles. The evidence BAMs do not contain reads and mappings for loci that were ultimately no-called or called homozygous ref, unless those regions were selected for de novo assembly because they were suspected to contain a variation. Every read that is found in the evidence files will also be present in the initial mappings, but the mapping positions may be different. In this case, where a read is found in both the *_mapping_sorted_header.bam and evidenceDnbs* files, the mapping in the evidence files is preferred.

### Complete Genomics LFR

The Complete Genomics LFR data was sequenced from cells sourced from Coriell. VCF and var formats files that include small variant calls are available for GM12878 and the Ashkenazi trio as indicated in Table 4. Both file formats include phasing information; see http://www.completegenomics.com/documents/DataFileFormats_Standard_Pipeline_2.5.pdf for details on file formats. In addition, summary files are included with assembly statistics. The VCF and var files also include two additional FORMAT fields: MEWC and SWC. MEWC, minimum exclusive well count, indicates the number of LFR wells that support the REF or ALT allele (whichever is fewer) exclusively, and not the other allele. SWC, shared well count, indicates the number of LFR wells that support both the REF and ALT allele. High confidence variant calls should have a high MEWC (typically greater than 3) and a low (ideally 0) SWC. Note that small variant sensitivity is somewhat lower for the LFR process compared to standard Complete Genomics sequencing, so the standard sequencing should be deferred to for unphased variants.

**Table 4:** Complete Genomics LFR data available at ftp://ftp-trace.ncbi.nlm.nih.gov/giab/ftp/

|  |  |  |
| --- | --- | --- |
| GM12878<br>(rep 1) | VCF, var<br>summary | /data/NA12878/CompleteGenomics_LFR/*GS000039392-ASM-NA12878* |
| GM12878<br>(rep 2) | VCF, var<br>summary | /data/NA12878/CompleteGenomics_LFR/*GS000039396-ASM-NA12878* |
| GM12878<br>(rep 3) | VCF, var<br>summary | /data/NA12878/CompleteGenomics_LFR/*GS000039473-ASM-NA12878* |
| GM24385 | VCF, var<br>summary | /data/AshkenazimTrio/HG002_NA24385_son/CompleteGenomics_LFR/*GS000039526-ASM-NA24385* |
| GM24143 | VCF, var<br>summary | /data/AshkenazimTrio/HG004_NA24143_mother/CompleteGenomics_LFR/*GS000039524-ASM-NA24143* |
| GM24149 | VCF, var<br>summary | /data/AshkenazimTrio/HG003_NA24149_father/CompleteGenomics_LFR/*GS000039541-ASM-NA24149* |

### Ion exome sequencing

The files generated by Thermo Fisher Scientific describe genomic variants called from Ion Torrent sequencing data with AmpilSeq exomes. The variants are represented in VCF files, each accompanied by an effective region BED file describing the corresponding genomic scope of called variants.

Four GIAB samples with were sequenced using AmpliSeq exome and sequenced on the Ion Proton (see https://www.lifetechnologies.com/order/catalog/product/4487084).

AmpliseqExome.20141120.16runs.vcf.zip -- 16 VCF files produced by Torrent Variant Caller v4.4 on 16 AmpliSeqExome runs, 4 of each samples picked by Genome in a Bottle consortium (NA24143, NA24149, NA24385 and NA24631).

AmpliseqExome.20141120.NA24143.vcf -- NA24143 variants called on 4 runs combined, above a quality score of 25;

AmpliseqExome.20141120.NA24149.vcf -- NA24149 variants called on 4 runs combined, above a quality score of 25;

AmpliseqExome.20141120.NA24385.vcf -- NA24385 variants called on 4 runs combined, above a quality score of 25;

AmpliseqExome.20141120.NA24631.vcf -- NA24631 variants called on 4 runs combined, above a quality score of 25;

AmpliseqExome.20141120_effective_regions.bed -- Genomic scope of AmpliSeqExome variant calls. This file describes the region in which variants are called with Torrent Suite v4.4 and later, and the region on which curation has been performed.

High_Confidence_Variants_NA24385.bed -- A list of inspected NA24385 variants based on Ion Torrent, Complete Genomics, 23andme and manual curation

High_Confidence_Variants_NA24385_effective_regions.bed -- Genomic scope of inspected NA24385 variants.

The Ion Exome data are available on the NCBI SRA [Data Citation 9] and [Data Citation 10], and on the GIAB FTP site at

ftp://ftp-trace.ncbi.nlm.nih.gov/giab/ftp/data/AshkenazimTrio/HG002_NA24385_son/ion_exome/

ftp://ftp-trace.ncbi.nlm.nih.gov/giab/ftp/data/AshkenazimTrio/HG003_NA24149_father/ion_exome/

ftp://ftp-trace.ncbi.nlm.nih.gov/giab/ftp/data/AshkenazimTrio/HG004_NA24143_mother/ion_exome/

ftp://ftp-trace.ncbi.nlm.nih.gov/giab/ftp/data/ChineseTrio/HG005_NA24631_son/ion_exome/

ftp://ftp-trace.ncbi.nlm.nih.gov/giab/ftp/data/NA12878/ion_exome/

### SOLiD 5500xl Wildfire WGS

The SOLiD 5500xl Wildfire WGS data is currently available as xsq files on the GIAB ftp site because this is the native format for SOLiD, as well as bam files mapped with lifescope and read groups for each lane. These data are available on the NCBI SRA [Data Citation 11], and here:

ftp://ftp-trace.ncbi.nlm.nih.gov/giab/ftp/data/AshkenazimTrio/HG002_NA24385_son/NIST_SOLiD5500W

ftp://ftp-trace.ncbi.nlm.nih.gov/giab/ftp/data/ChineseTrio/HG005_NA24631_son/NIST_SOLiD5500W

### Bionano Genomics genome maps

all.bnx is the raw data after image processing and filtering for molecules >150kb

EXP_REFINEFINAL1.cmap is the de novo assembly consensus genome map set

The following files result from the alignment of genome maps to hg19:

EXP_REFINEFINAL1.xmap is the alignment file with match group information

EXP_REFINEFINAL1_q.cmap is the de novo genome maps that align to hg19 (query, it’s a subset of all genome maps)

EXP_REFINEFINAL1_r.cmap is an in silico map of hg19 (Nt.BspQI motifs, anchor in the alignment)

BioNano data are available at

ftp://ftp-trace.ncbi.nlm.nih.gov/giab/ftp/data/AshkenazimTrio/HG002_NA24385_son/BioNano/

ftp://ftp-trace.ncbi.nlm.nih.gov/giab/ftp/data/AshkenazimTrio/HG003_NA24149_father/BioNano/

ftp://ftp-trace.ncbi.nlm.nih.gov/giab/ftp/data/AshkenazimTrio/HG004_NA24143_mother/BioNano/

ftp://ftp-trace.ncbi.nlm.nih.gov/giab/ftp/data/ChineseTrio/HG005_NA24631_son/BioNano/

### PacBio

The PacBio data are available on the NCBI SRA [Data Citation 12] and on the GIAB FTP site at:

ftp://ftp-trace.ncbi.nlm.nih.gov/giab/ftp/data/AshkenazimTrio/HG002_NA24385_son/PacBio_MtSinai_NIST/

ftp://ftp-trace.ncbi.nlm.nih.gov/giab/ftp/data/AshkenazimTrio/HG003_NA24149_father/PacBio_MtSinai_NIST/

ftp://ftp-trace.ncbi.nlm.nih.gov/giab/ftp/data/AshkenazimTrio/HG004_NA24143_mother/PacBio_MtSinai_NIST/

The file/directory naming convention is defined as follows:

[SampleName]/[WellName]_[CollectionNumber].[UUID].tar.gz Note that SampleName may contain other genomes in the name since this is hardcoded by the run name, but the data directories only contain run data from AJ son, AJ father, and AJ mother. For example, for SampleName of HG002new_O1_BP_P6_021815_MB_105pM, WellName of A01, and CollectionNumber of 3, you will see a tar.gz file in HG002new_O1_BP_P6_021815_MB_105pM directory with name A01_3.[UUID].tar.gz The UUID is currently used for only hashing purpose. The tar.gz file contains the raw SMRTPortal data including following contents:

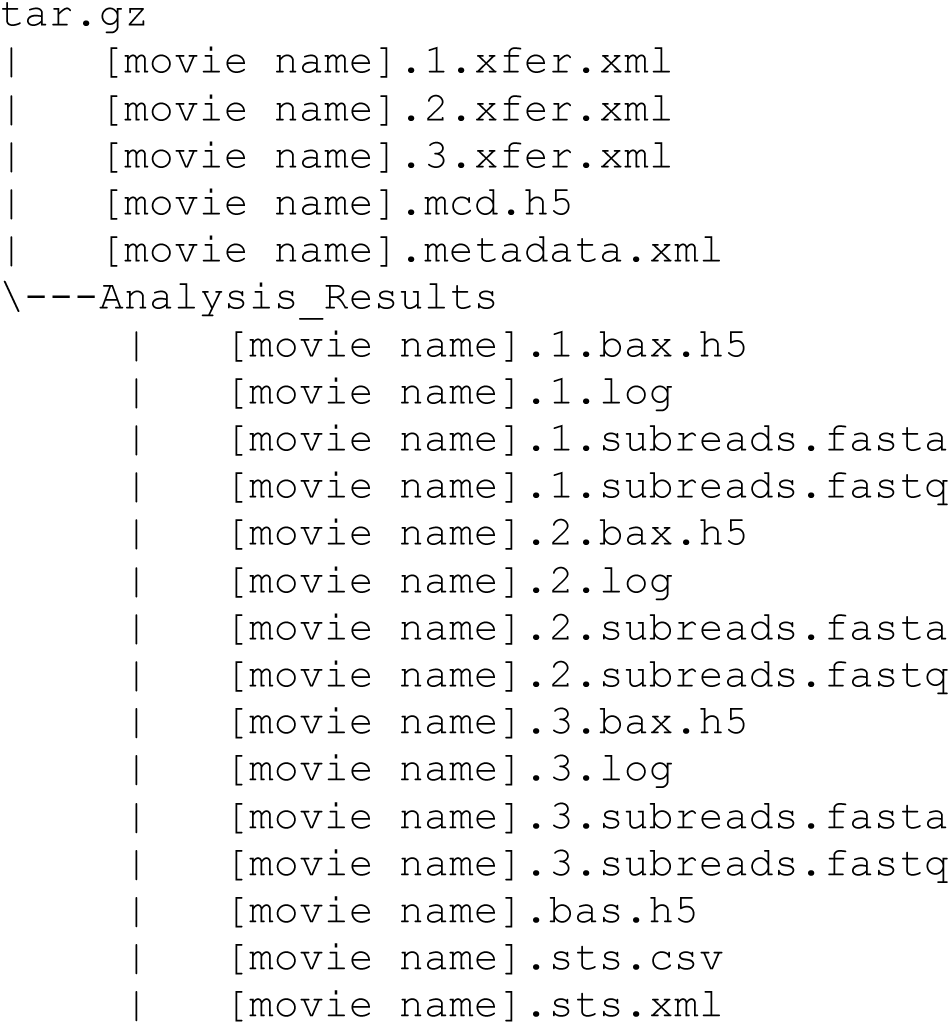

The metadata.xml contains all the metadata of this particular sample in the xml format; for example, in the TemplatePrep field you might see “DNA Template Prep Kit 2.0 (3Kb - 10Kb),” and in the BindingKit field you might see “DNA/Polymerase Binding Kit P6,” etc. For information about bas.h5/bax.h5 files, please see:

http://files.pacb.com/software/instrument/2.0.0/bas.h5%20Reference%20Guide.pdf

For information about subreads, please see: https://speakerdeck.com/pacbio/track-1-de-novo-assembly

### Oxford Nanopore

The Oxford Nanopore raw reads and 2D reads are available at:

ftp://ftp-trace.ncbi.nlm.nih.gov/giab/ftp/data/AshkenazimTrio/HG002_NA24385_son/CORNELL_Oxford_Nanopore/

### Future Data and FTP structure

We expect to continue to accrue public data for these genomes as new methods become available. These data will be placed in the NCBI SRA when possible, linked to the GIAB BioProject PRJNA200694 and the appropriate BioSample listed in Table 2. Other data and analyses will also be publically available on the GIAB FTP site at NCBI (ftp://ftp-trace.ncbi.nih.gov/giab/ftp). Preliminary data will be placed in the technical directory and analyses and finalized data will be placed under each trio or genome in the data directory. The directories under the analysis directory for each family contains the institution, data set, type(s) of variants, analysis tool, and date.

## Technical Validation

### Illumina paired end WGS

Several statistics were calculated for each flow cell using the Illumina BaseSpace Isaac Whole Genome Sequencing v3 analysis pipeline (Table 5).

**Table 5:** Illumina paired end sequencing statistics from Illumina BaseSpace Isaac Whole Genome Sequencing v3 analysis pipeline *(summarized by genome)*

| Genome | Cov-<br>erage<br>(x) | Read<br>length | percent<br>duplicate<br>paired<br>reads | Fragment<br>Length<br>Median<br>(bp) | Fragment<br>length<br>standard<br>deviation<br>(bp) | read 1<br>percent<br>aligned | read 2<br>percent<br>aligned | read 1<br>mismatch<br>rate (%) | read 2<br>mismatch<br>rate (%) |
| --- | --- | --- | --- | --- | --- | --- | --- | --- | --- |
| HG002 | 290 | 2x150 | 1.3 | 567 | 150 | 95.6 | 94.2 | 0.49 | 0.73 |
| HG003 | 294 | 2x150 | 1.3 | 562 | 146 | 94.8 | 93.2 | 0.49 | 0.73 |
| HG004 | 324 | 2x150 | 1.3 | 562 | 143 | 96.0 | 93.8 | 0.49 | 0.84 |
| HG005 | 306 | 2x250 | 3.9 | 577 | 154 | 96.7 | 94.6 | 0.97 | 1.58 |
| NA24694 | 105 | 2x150 | 3.7 | 560 | 146 | 95.8 | 93.9 | 0.51 | 0.78 |
| NA24695 | 106 | 2x150 | 3.5 | 560 | 140 | 96.2 | 93.9 | 0.49 | 0.83 |

### Illumina mate-pair WGS

To assess duplication rate, coverage, and insert size of the mate-pair libraries, reads were stripped of adapter sequences. Read pairs were removed if the sequence of one or both mates was less than 20 bp after adapter stripping, or if the adapter sequence was at the beginning rather the end of a read (indicating the read inserts were likely to be in inward-facing F/R orientation rather than the expected outward-facing R/F orientation). Reads were then mapped to the hg19 reference genome using “bwa mem” (Li 2013) with default settings, and duplicates were marked using samblaster (Faust 2014). Statistics are summarized in Table 6.

**Table 6:** Mate-pair sequencing statistics.

|  | after adapter stripping |  |  |  | sequence coverage |  | physical coverage |  | insert size |  |
| --- | --- | --- | --- | --- | --- | --- | --- | --- | --- | --- |
|  | read count | base count | uniquely mapping (mapq=60, | percent dups | mean | std | mean | std | mean | std |

|  |  | (Gb) | removing dups) |  |  |  |  |  |  |  |
| --- | --- | --- | --- | --- | --- | --- | --- | --- | --- | --- |
| **HG002** | 890,861,081 | 94.5 | 366,322,711 | 51 | 13.6 | 4.8 | 447 | 48.8 | 6466 | 1220 |
| **HG003** | 822,621,264 | 89.5 | 353,618,533 | 49 | 13.6 | 4.7 | 408 | 45.4 | 6110 | 934 |
| **HG004** | 815,116,039 | 88.5 | 379,255,921 | 45 | 14.2 | 4.9 | 421 | 49.4 | 6052 | 1375 |
| **HG005** | 663,052,149 | 71.4 | 382,392,628 | 32 | 14.3 | 5.1 | 463 | 51.7 | 6290 | 1982 |
| **NA24694** | 663,071,189 | 71.9 | 436,995,349 | 22 | 16.7 | 5.7 | 525 | 60.2 | 6517 | 1149 |
| **NA24695** | 648,828,312 | 71.1 | 418,948,777 | 24 | 15.8 | 5.5 | 507 | 64.9 | 6721 | 1204 |

The high rate of PCR duplicates (close to 50% in some libraries) resulted in lower than expected sequence coverage (13–17x average across all sequenced genomic positions). A more relevant metric for mate-pair data is the physical coverage, which measures the number of inferred fragments that cover a particular genomic position (including both the sequenced ends as well as the unsequenced genomic region between the ends). Because the empirical insert size average was between 6–7kb per individual, the physical coverage of the genome was quite high (>400x per individual). BAMs were stripped of duplicate reads to reduce file size, but the full data are available in fastq format.

### Illumina read clouds (synthetic long reads)

Several statistics were generated to assess the Illumina read clouds (Figure 2): the read coverage distribution for each cloud, the fragment coverage distribution for the whole genome, the distribution for cloud length, and the probability density estimation for template length (same over samples). The cloud length distributions are plotted by type of the clouds. In such figures, ‘0 end’ means both of the end-markers of a cloud are missing, and so on. Thus for clouds with both end-markers (2 ends), the expectation for the length is larger. Also for genome coverage distribution, the y-axis is the length of the genome covered by corresponding fragment coverage (integrate to 3Gb).

### Illumina paired end WES

To assess duplication rate, coverage, and insert size of the libraries, picard CollectAlignmentSummaryMetrics, CollectInsertSizeMetrics and CalculateHsMetrics were performed on each sample BAM file. Statistics are summarized in Table 7.

**Table 7:** Illumina paired end WES library statistics using Picard

|  | HG002 | HG003 | HG004 | HG005 |
| --- | --- | --- | --- | --- |
| % Bases in the fastq files which have base quality higher than 30 | 93.0 | 93.0 | 92.9 | 91.7 |
| Mean target coverage | 225 | 192 | 213 | 181 |
| Mean insert size | 201 | 203 | 200 | 203 |
| % Duplication | 4.9 | 4.2 | 4.9 | 5.7 |
| % Pass Illumina filter reads aligned to the human reference genome | 99.7 | 99.7 | 99.7 | 99.8 |
| % Target bases covered by 20 reads or more | 98.6 | 98.4 | 98.6 | 98.3 |
| % Target bases covered by 30 reads or more | 97.9 | 97.5 | 97.9 | 97.2 |

### 10X Genomics GemCode™ Libraries for Illumina Sequencing

Statistics for the AJ trio and Chinese son were generated using the 10X GemCode Long Ranger software (Table 8).

**Table 8:**
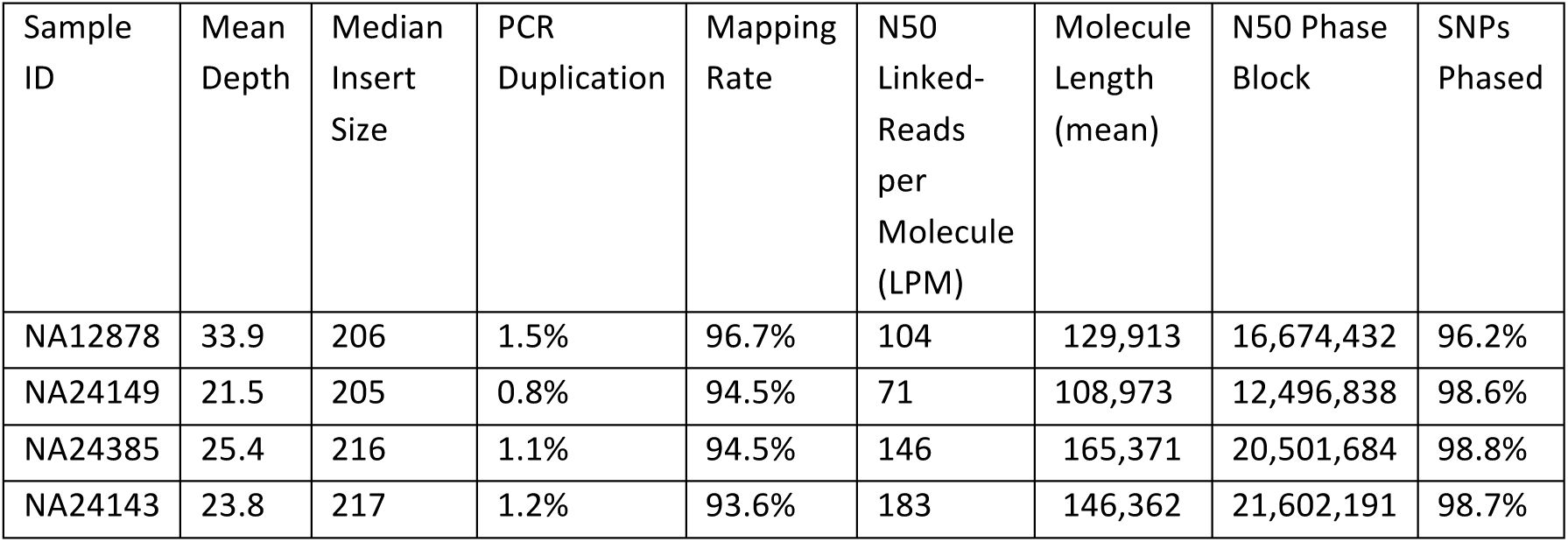
10X Genomics library statistics generated using the GemCode Long Ranger software package.

| Sample ID | Mean Depth | Median Insert Size | PCR Duplication | Mapping Rate | N50 Linked-Reads per Molecule (LPM) | Molecule Length (mean) | N50 Phase Block | SNPs Phased |
| --- | --- | --- | --- | --- | --- | --- | --- | --- |
| NA12878 | 33.9 | 206 | 1.5% | 96.7% | 104 | 129,913 | 16,674,432 | 96.2% |
| NA24149 | 21.5 | 205 | 0.8% | 94.5% | 71 | 108,973 | 12,496,838 | 98.6% |
| NA24385 | 25.4 | 216 | 1.1% | 94.5% | 146 | 165,371 | 20,501,684 | 98.8% |
| NA24143 | 23.8 | 217 | 1.2% | 93.6% | 183 | 146,362 | 21,602,191 | 98.7% |

### Complete Genomics WGS

#### Genomic Assembly

Sequencing results in mate-paired reads with a 2-4 base overlap between adjacent contiguous sequences, as described in the “Read Data Format” section of the Data File Formats documentation (http://www.completegenomics.com/documents/DataFileFormats_Standard_Pipeline_2.5.pdf). The gapped read pairs were aligned to the NCBI Build 37 reference gnome using an index lookup based fast algorithm. At locations where the mapping results suggest the presence of a variant, mapped reads were refined, expanded and then assembled into a best-fit, diploid sequence with a custom software suite employing both Bayesian and de Bruijn graph techniques.^9^ This process yielded diploid reference, variant or no-call at each genomic location with associated variant quality scores. In addition to small variants, larger variants are detected, including MEIs, CNVs and SVs. The Complete Genomics CNV pipeline has the following steps: 1) Various measures of coverage for tiled 2kb and 100kb windows across the genome are determined. This is provided in the cnvDetails* and depthOfCoverage files. 2) The genome is segmented into called ploidy levels (diploid model, 2kb windows) or coverage levels (non diploid model, 100 kb windows) using diploid and non-diploid HMM-based algorithms. The segmentation patterns called are provided in the cnvSegments* files. 3) The lesser allele fraction (LAF) is calculated for 100 kb windows across the genome - the LAF calculations are included in the cnvDetailsNondiploid and cnvSegmentsNondiploid files (and not in the diploid files because of the 100kb window size restriction). To identify SVs, DNB mappings found during the standard assembly process are analyzed to find clusters of DNBs in which each arm maps uniquely to the reference genome, but with an unexpected mate pair length or anomalous orientation. SVs are encoded in the junctions and highConfidenceJunctions files where the latter file contains a high-confidence filtered subset of the data in former file. In addition to calling structural variant junctions, junctions are rationalized into structural variation events using the CGA Tools junctions2events algorithm. These data are provided in the svEvents and highConfidenceSvEvents files. Additional information on the CNV, SV and MEI algorithms is available here: http://www.completegenomics.com/documents/DataFileFormats_Standard_Pipeline_2.5.pdf Assembly metrics are summarized in Table 9. Additional summary information can be found for each genome in the full package in the ASM/summary-*.tsv file, see Table 3.

**Table 9:** Complete Genomics WGS Summary Metrics

| Subject | GM12878 | GM24385 | GM24143 | GM24149 | GM24631 | GM24631<br>(Coriell<br>cells) | GM24695<br>(Coriell<br>cells) | GM24694<br>(Coriell cells) |
| --- | --- | --- | --- | --- | --- | --- | --- | --- |
| Fully called genome fraction | 0.975 | 0.977 | 0.975 | 0.976 | 0.976 | 0.976 | 0.974 | 0.976 |
| Gross mapping yield (Gb) | 341 | 360 | 376 | 350 | 356 | 353 | 352 | 355 |
| Both mates mapped yield (Gb) | 313 | 322 | 351 | 313 | 320 | 318 | 314 | 318 |
| Mate distribution mean | 389 | 395 | 405 | 386 | 395 | 374 | 379 | 387 |
| SNP total count (PASS only) | 3449567 | 3463164 | 3501307 | 3436943 | 3431805 | 3424251 | 3445736 | 3407542 |
| SNP transitions/transversions ratio (PASS only) | 2.125 | 2.127 | 2.122 | 2.128 | 2.126 | 2.128 | 2.125 | 2.127 |

### Complete Genomics LFR

#### Genomic Assembly

Genomic assembly was performed as described above for Regular Complete Genomics WGS, with the added assembly step of haplotype generation using well information.^8^ LFR assemblies include only small variants and their associated haplotypes and well counts.

Assembly metrics are summarized in Table 10. Additional data summary information can be found for each genome in the full package in the ASM/summary-*.tsv file, see Table 4.

**Table 10:** Complete Genomics LFR WGS Summary Metrics

| Subject | GM12878 (rep1) | GM12878 (rep2) | GM12878 (rep3) | GM24385 | GM24143 | GM24149 |
| --- | --- | --- | --- | --- | --- | --- |
| Fully called genome fraction | 0.952 | 0.968 | 0.971 | 0.972 | 0.968 | 0.972 |
| Gross mapping yield (Gb) | 363 | 362 | 368 | 365 | 371 | 363 |
| Both mates mapped yield (Gb) | 317 | 311 | 316 | 310 | 325 | 312 |
| Mate distribution mean | 310 | 290 | 290 | 275 | 277 | 282 |
| SNP total count (PASS only) | 3308426 | 3410795 | 3428944 | 3443940 | 3461563 | 3418049 |
| SNP transitions/transversions ratio (PASS only) | 2.134 | 2.134 | 2.134 | 2.135 | 2.134 | 2.140 |
| N50 | 181kb | 257kb | 176kb | 887kb | 1661kb | 734kb |
| Median Contig Length | 76kb | 84kb | 74kb | 133kb | 149kb | 125kb |
| Fraction of Phased SNPs | 98.7% | 99.4% | 99.4% | 99.7% | 99.6% | 99.6% |

### Ion exome sequencing

Sequencing reads with a mean read length of 190bp were mapped to human genome version hg19. Mean coverage across AmpliSeq™ Exome target regions is 256x per run, with raw read accuracy at 99%. Additional summary statistics are in Table 1.

### SOLiD 5500xl Wildfire WGS

As recommended by the manufacturer, the statistics reported from the instrument from each run of the SOLiD 5500xl Wildfire WGS were examined to ensure consistency in quality. These statistics included “Quality Value” and “Fraction of good+best”, which are a function of the quality of the signal at each ligation. Reads were mapped using Lifescope, which generated the summary statistics given in Table 11.

**Table 11:**
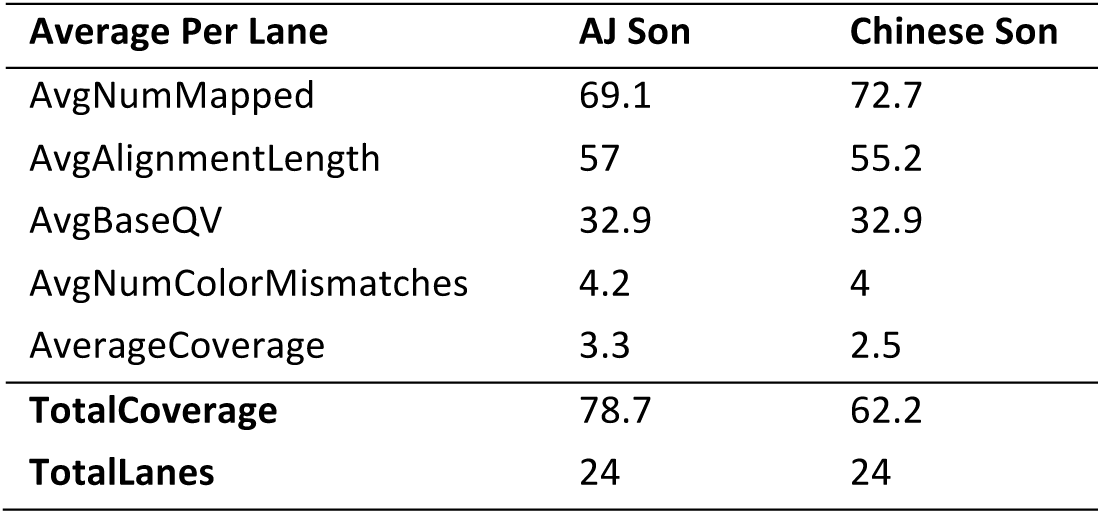
SOLiD 5500xl Wildfire WGS statistics from Lifescope mapped reads

### Bionano Genomics genome maps

De novo assembly of single molecules is accomplished using BioNano Genomics IrysSolve™, a proprietary assembler software application, based on an overlap-layout-consensus paradigm.^10-12^ Molecules longer than 150 kb were the input for a pairwise comparison to find all overlaps; then a draft consensus map (BioNano Genomics CMAP) was constructed based on these overlaps. The draft BioNano Genomics CMAP was refined by mapping single molecules to it and iteratively recalculating the label positions. Next, the draft BioNano Genomics CMAP (consensus genome maps) were extended by aligning overhanging molecules to the consensus maps and calculating a consensus in the extended regions. Finally, the consensus maps were compared and merged iteratively five times where the patterns matched and then the final label position calculation was made. Summary statistics for each sample are presented in Table 12.

**Table 12:**
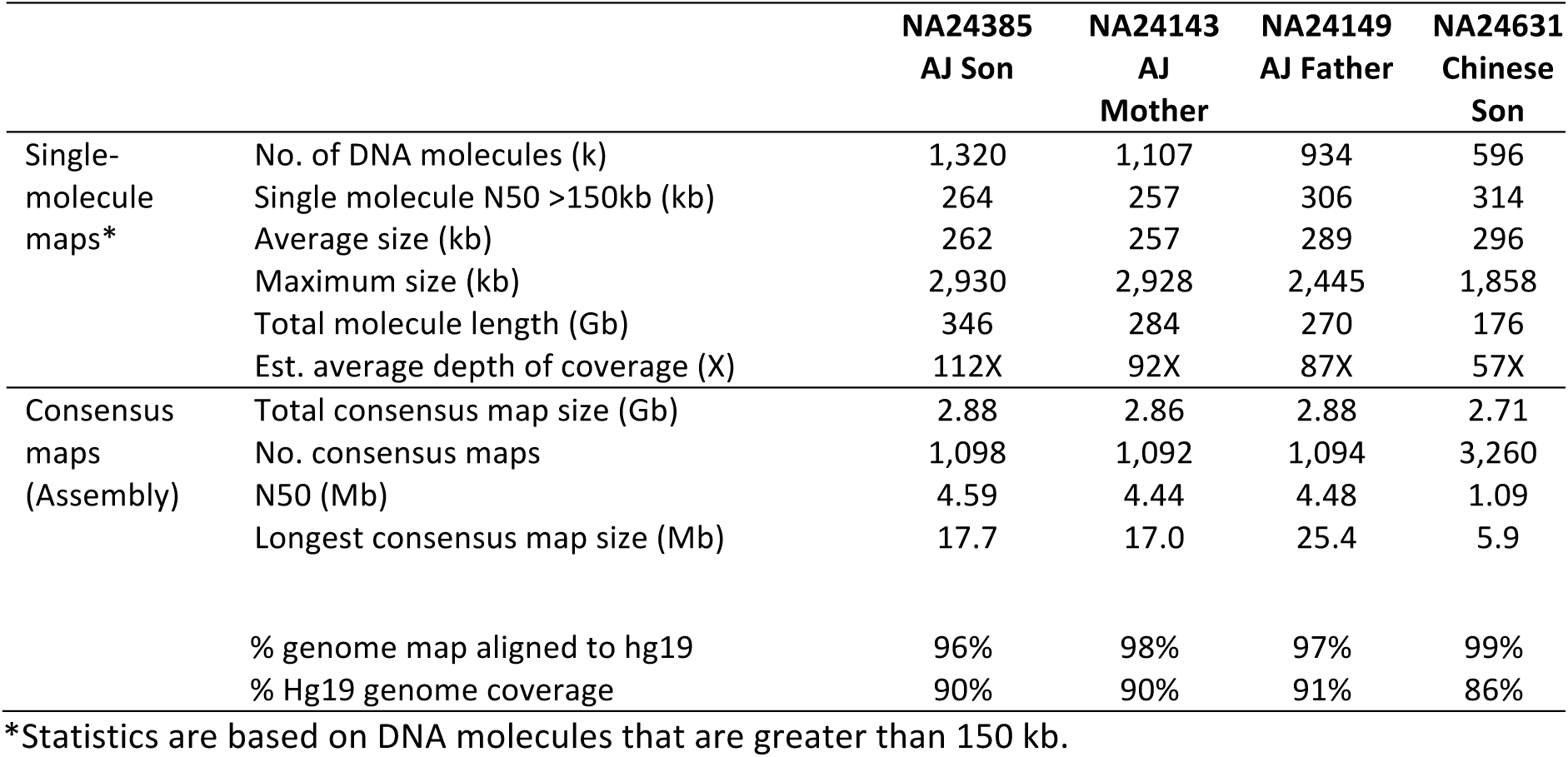
Bionano statistics for mapping of single molecules as well as the assembly “consensus maps”

### PacBio

Assuming a 3.2 Gb human genome, sequencing was conducted to approximately 69X, 32X, and 30X coverage for AJ son, AJ father, and AJ mother across 292, 139, and 132 SMRT cells, respectively. 27.4M, 13.2M, and 12.4M subreads were generated resulting in 220.0, 101.6, and 94.9 Gb of sequence data with sub-read length N50 values of 11,087, 10,728, and 10,629 basepairs. The coverage distribution for each genome is also depicted in Figure 3.

### Oxford Nanopore

Data from two libraries from Oxford Nanopore giving about 0.017x coverage is described in ^14^.

## Usage Notes

The genomes sequenced in this work (Table 2) and their data (Table 1) are all publicly available both as cell lines and as DNA. The pilot genome, NIST RM 8398 (based on Coriell DNA NA12878), is available both from Coriell as well as from NIST (http://tinyurl.com/giabpilot). The NIST RM 8398 was prepared by Coriell from a large growth of cells, and the DNA was extracted and mixed to produce about 8300 10 ug vials of DNA. The remaining genomes are from the Personal Genome Project. These genomes are also available as EBV-immortalized B lymphoblastoid cell lines and as extracted DNA from Coriell, and 4 of them will be available as NIST RMs, planned for release in early 2016. The AJ Son will be distributed as RM 8391, the AJ Trio will be distributed as RM 8392, and the Chinese Son will be distributed as RM 8393 (note that the Chinese parents are only available from Coriell). Similar to the pilot genome, the other candidate NIST RMs are extracted DNA from a large batch of cells. Except for technologies that optimally start with cells (Complete Genomics LFR, 10X Genomics, and BioNano), all data in this work are collected from the NIST RM DNA. It is possible that small differences may exist between the NIST RM DNA and the DNA from Coriell because they come from different passages of cells and may contain different new mutations.

All data from Genome in a Bottle project are available without embargo, and the primary location for data access is ftp://ftp-trace.ncbi.nlm.nih.gov/giab/ftp. To facilitate data analysis in cloud, all the data have been mirrored to the Amazon cloud with “s3://giab” as bucket name. In addition, data that were submitted to SRA can also be accessed through NCBI BioProject (http://www.ncbi.nlm.nih.gov/bioproject/200694). The Genome in a Bottle Consortium has formed an Analysis Team to coordinate analyses by groups that are interested in analyzing these data. The primary goal of this group is to establish high-confidence phased variant calls of all sizes for these genomes, so that anyone can benchmark accuracy of their calls for these genomes. The Analysis Team has several sub-groups working on assembly, small variant calling, structural variant calling, and phasing. The intermediate analysis results from these sub-groups are being organized in subdirectory under “analysis” (ftp://ftp-trace.ncbi.nlm.nih.gov/giab/ftp/data/{Ashkenazim|Chinese}Trio/analysis/) with the name describing analyzer’s name who performed the analysis, technology for dataset(s) that has been used, type of variant being characterized, analysis tool or algorithm being utilized, and the submission date (MMDDYYYY format) serving as version for better understanding what the datasets were about. The integrated high-confidence calls for the trio samples will be available at ftp://ftp-trace.ncbi.nlm.nih.gov/giab/ftp/release/, and the subdirectory with name “latest” will always contain the latest results published by the Genome in a Bottle Consortium. To make it easier to visualize these data in a web-based genome browser, the GeT-RM browser at NCBI is also hosting some of the vcf and bam files from NA12878 and is currently working on hosting additional data from NA12878 as well as the other GIAB genomes.

In order to improve the accessibility and usability of the Genome in a Bottle project data, a GitHub site (https://github.com/genome-in-a-bottle) has also been developed for the Genome in a Bottle project. The idea behind this site is to help the users navigate the main ftp site more easily, and find as much answers as possible by themselves related to the Genome in a Bottle project and its data. Several repositories have been established within the genome-in-a-bottle GitHub site. The repository of “about_GIAB” describes the objectives and overview of the Genome in a Bottle project, while the repository of “giab_data_indexes” lists and organizes the index files for the raw sequences and/or alignments by the sequencing platforms for each of the individuals or trio families, therefore, users could use the specific index file to guide their downloading for the desired dataset from the ftp site. The repositories of “giab_data_analysis” and “giab_latest_release” provide easy links to the specific ftp locations regarding ongoing analysis results performed by individual analysis groups and the latest high-confidence sets released by the Genome in a Bottle Consortium. The tools and methods that have been used by the analysis groups have been documented in “giab_tools_methods” repository, while the scientific publications are listed in “giab_publications” repository. The “giab_FAQ” repository provides short answers for the frequently asked questions regarding how to effectively access and appropriately utilize the data from the Genome in a Bottle project.

## Acknowledgements

We thank members of the Genome in a Bottle Consortium for their feedback before and during this study. Certain commercial equipment, instruments, or materials are identified in this paper only to specify the experimental procedure adequately. Such identification is not intended to imply recommendation or endorsement by the NIST, nor is it intended to imply that the materials or equipment identified are necessarily the best available for the purpose.

## Author contributions

JMZ, MLS, and GIAB designed the overall study.

JMZ, DC, JM, LV, NS, ZW, YL, NA, EH, EJ, RS, RMT, KZ, YF, MC, CX, and MLS wrote the manuscript.

JMZ, DC, JM, LV, and MLS designed, sequenced, and analyzed the Illumina paired end WGS.

JMZ, DC, JM, NS, AS, ZW, and MLS designed, sequenced, and analyzed the Illumina mate pair WGS.

JMZ, DC, JM, NS, AS, YL, FC, EJ, AM, and MLS designed, sequenced, and analyzed the Illumina synthetic long read WGS.

CM, NA, and EH designed, sequenced, and analyzed the Oxford Nanopore WGS.

KP, WS, TL, MS, ZD, AH, and HC designed, sequenced, and analyzed the BioNano mapping.

RMT, CCC, and NG designed, sequenced, and analyzed the Complete Genomics WGS.

KZ, SG, FH, and YF designed, sequenced, and analyzed the Ion Torrent exome sequencing.

JMZ, JM, GD, ES, RS, AB, MC, and MLS designed, sequenced, and analyzed the PacBio WGS.

JMZ, AWZ, MB, JB, PE, GMC, MLS, and GIAB designed the process for selecting the genomes from the PGP

PM, SK-P, GSYZ, MS-L, HSO, and PAM designed, sequenced, and analyzed the 10X Genomics data

YS and KBR designed, sequenced, and analyzed the Illumina WES data

JMZ, SS, JT, and CX designed and manage the GIAB FTP site and SRA submissions

Bibliographic information for any works cited in the above sections, using the standard *Nature* referencing style.

## Data Citations

Bibliographic information for the data records described in the manuscript.

1. Zook, J.M., et al. NCBI SRA SRX1049768 to SRX1049855 (2015)
2. Zook, J.M., et al. NCBI SRA SRX847862 to SRX848317 (2015)
3. Zook, J.M., et al. NCBI SRA SRX1388368 to SRX1388459 (2015)
4. Zook, J.M., et al. NCBI SRA SRX1388732 to SRX1388743 (2015)
5. Sheng, Y., et al. NCBI SRA SRP047086 (2015).
6. Schnall-Levin, M., et al. NCBI SRA SRX1392293 to SRX1392296 (2015)
7. Truty, R., et al. NCBI SRA SRX840234 (2014)
8. Truty, R., et al. NCBI SRA SRX852932 to SRX852936 (2014)
9. Hyland, F., et al. NCBI SRA SRX847094 (2014)
10. Hyland, F., et al. NCBI SRA SRX848742 to SRX848744 (2014)
11. Zook, J.M., et al. NCBI SRA xxxx (2015)
12. Sebra, R., et al. NCBI SRA SRX1033793 to SRX1033798 (2015)

**Supplementary Figure 1:**
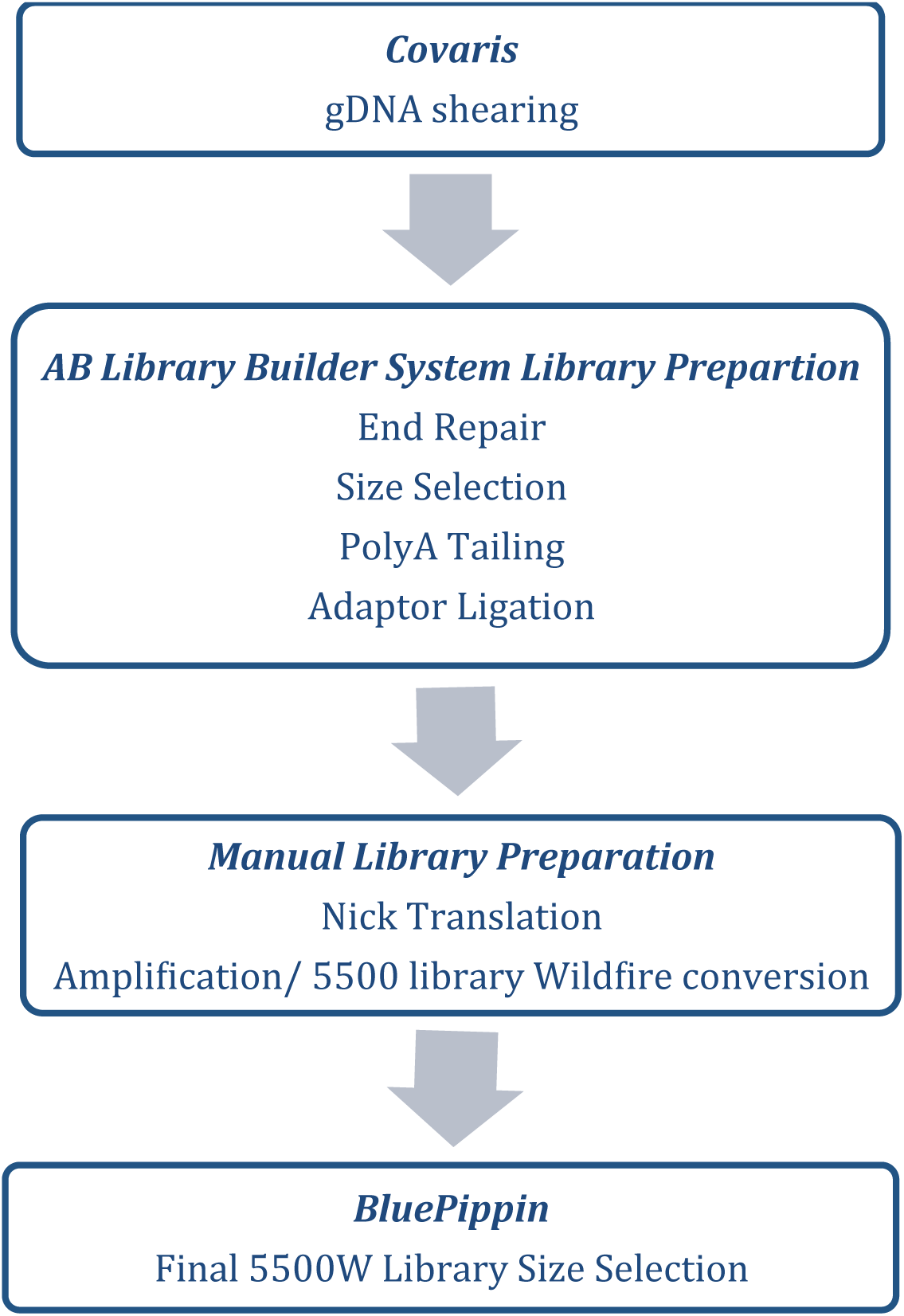
Workflow to produce libraries for 5500W sequencing of HG-005

**Supplementary Figure 2:**
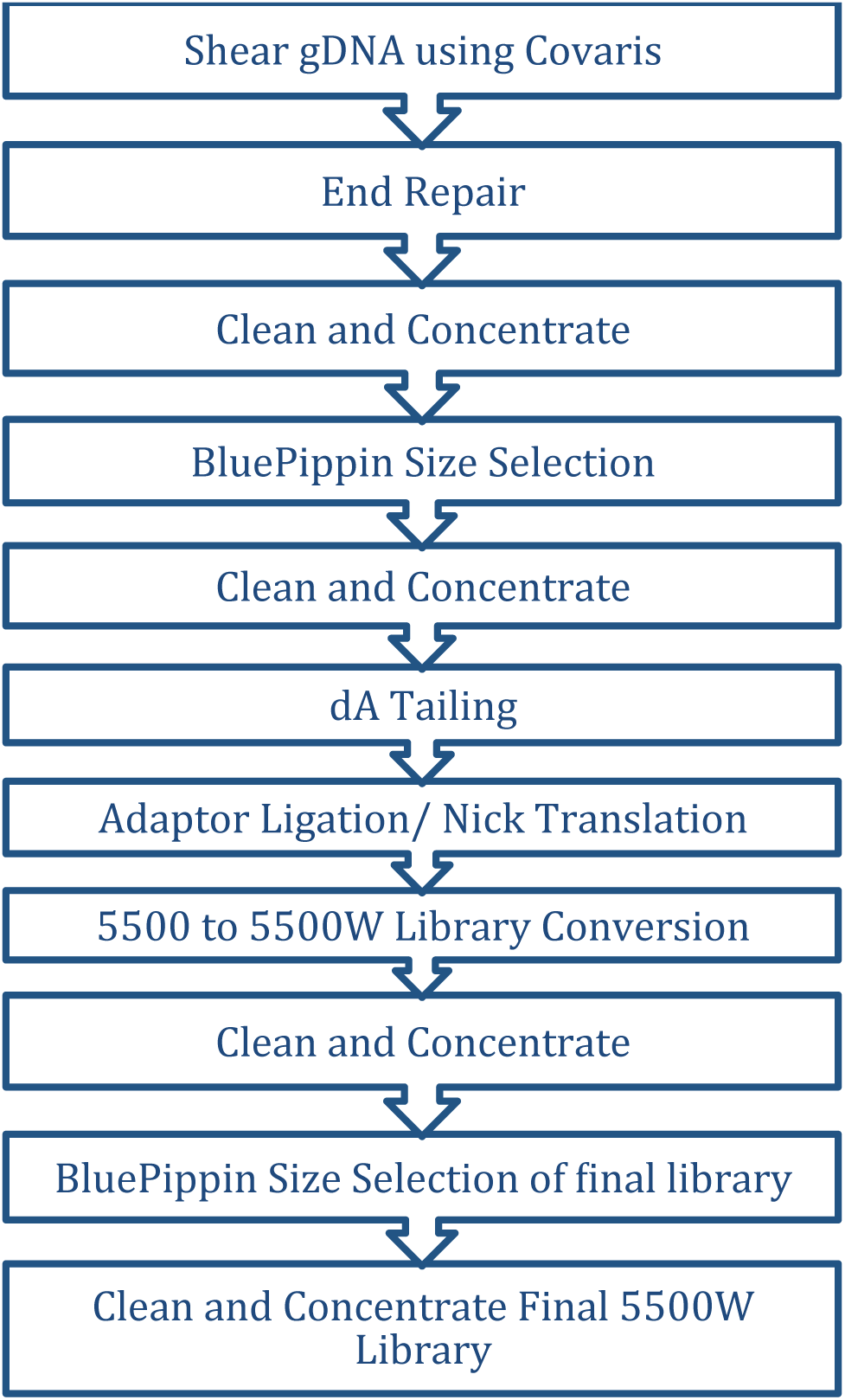
Workflow to produce libraries for 5500W sequencing of HG-002

**Supplementary Figure 3:**
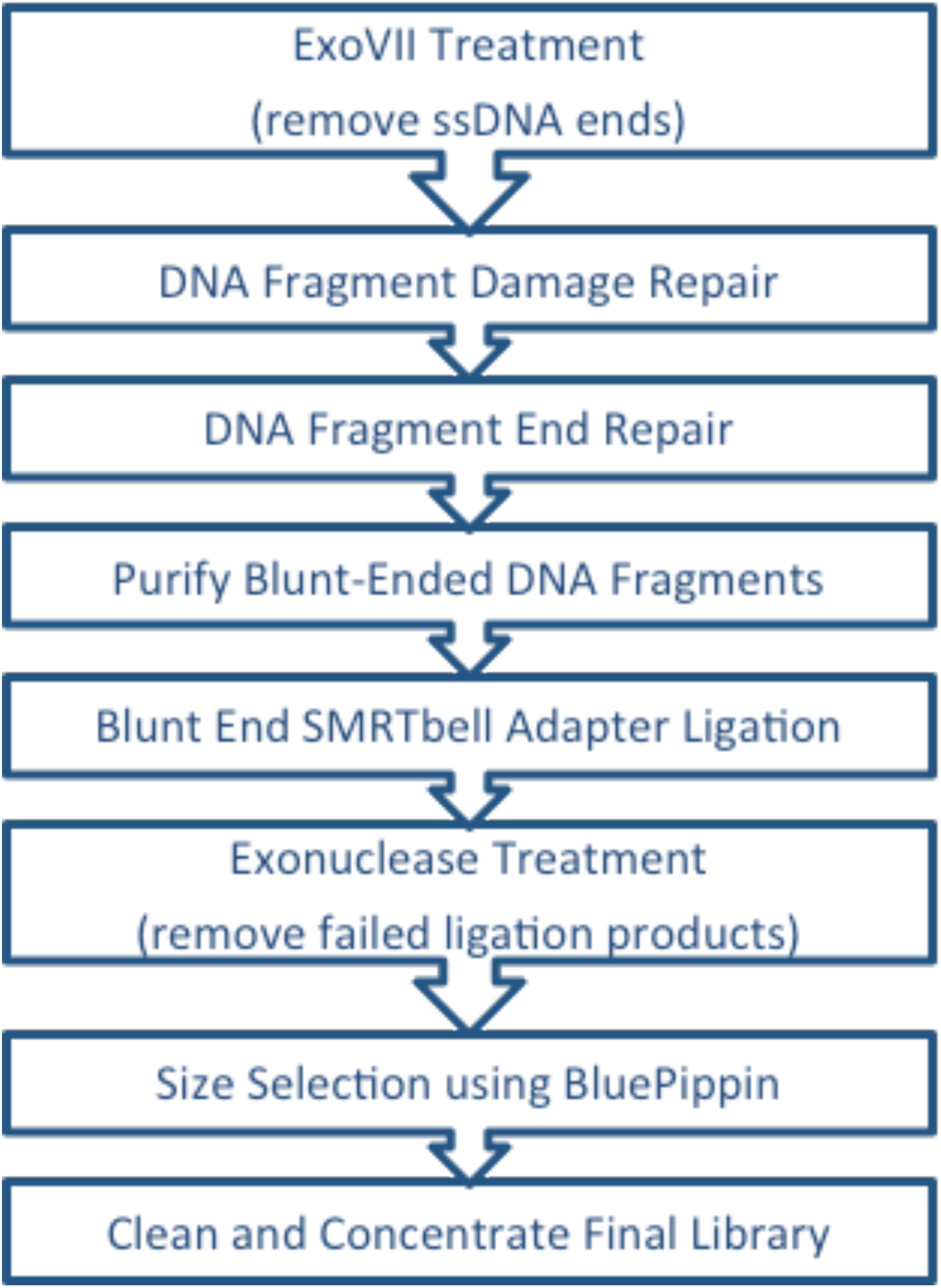
Workflow to produce libraries for Pacific Biosciences sequencing

